# High-throughput *in vivo* DNA data encoding by multiplexed conjugation and orthogonal base editing

**DOI:** 10.64898/2026.10.08.757515

**Authors:** Louise F.H. Funke, Zhangyuan Lin, Wenjun Liao, Yutong Ji, Chueh Loo Poh

## Abstract

Bacterial conjugation and CRISPR base editing offer complementary routes for programmable DNA transfer and heritable genomic writing, but their combined potential for information encoding in living cells remains unexplored. Here, we introduce CONCORD, a bacterial encoding platform that couples multiplexed bacterial conjugation to genomic base editing for synthesis-free *in vivo* DNA data writing. A recipient ‘Writer’ strain expresses the base-editing machinery and is equipped with a writing template, whereas reusable donor ‘Guide’ strains deliver the encoding sgRNA signals by conjugation. Using this setup, we encode binary information into genomic DNA in a high-throughput manner. Moreover, we systematically evaluated multiple orthogonal base editors for an expanded encoding repertoire and improved data recovery through redundancy encoding. We deploy the system to encode 45,689 bits into DNA including two culturally and historically significant artifacts: a greeting from NASA’s Voyager Golden Record and a photograph of Navajo petroglyphs and recovered these datasets with 100% and 99.5% accuracy, respectively, without computational error correction. Connecting and engineering the unique mating and immune system of microbial cells we have established a framework for programmable information writing in bacterial populations, with applications for biological event recording and sustainable DNA data storage.

## Introduction

Biological systems naturally record and store information, from hereditary information encoded in genomic DNA to adaptive and epigenetic memories that can influence future cellular behaviour^1,2^. Bacterial cells can preserve past encounters through stable genetic changes, including the acquisitions of CRISPR spacers and antibiotic resistance genes following encounters with viruses and other microbes, respectively^3,4^. These examples illustrate that biological systems are not merely passive carriers of information, but active, self-renewing, systems that write, copy and maintain new information in DNA.

Biological systems have been engineered to encode information for sensing and recording of biological events, responding to external signals such as chemicals, electrical stimuli or light^5–7^. Some engineered cellular information recorders could be applied to higher-level information encoding and were used for cellular lineage tracing^8^ or cellular computing^9^. Nevertheless, these in vivo information encoding systems have been constrained by limited writable capacity per cell due to inefficient encoding reactions and reliance on external inputs, resulting in low information encoding density and scalability^10^. In microbial systems, genome editing and horizontal gene transfer are particularly attractive components for scalable and high-throughput information encoding. CRISPR-guided base editors enable precise and efficient nucleotide conversion, making them well suited for the stable writing of binary states into DNA^11^. Bacterial conjugation, in turn, provides a simple and efficient mechanism for cell-to-cell DNA transfer that is initiated by mixing donor and recipient populations^12,13^. Despite the programmability of base editing and scalability of conjugative transfer, these two processes have not been combined for information writing in bacteria.

The global volume of digital information continues to increase, with a substantial fraction being “cold” data retained for archival rather than rapid access^14,15^. DNA is uniquely suited as a data storage medium because of its high information density^16–18^, low energy demand^17^, and long physical lifetime^19–21^, and particularly suitable for cold data storage due to its comparably slow information writing and reading processes. However, practical deployment has largely depended on de novo DNA synthesis, which, despite rapid advances, remains expensive, chemically intensive, and error-prone at scale^22–29^. Several de novo synthesis-free strategies have been explored for large-scale information encoding in DNA, including epigenetic bit printing^30^, DNA tape recording via base conversion^11^, and strand-nick-based DNA punch cards^31^. Although these methods reduce dependence on DNA synthesis, they still depend on synthesized DNA in the form of template strands and input materials, for example ssDNA bricks^30^ or sgRNA^11^. Moreover, they rely on enzymatic in vitro reactions, which require purified enzymes and chemicals as additional inputs^21^. A more sustainable alternative route is to encode information directly in living cells, where cells can generate the DNA substrate, express the molecular writing machinery and preserve the encoded information through genome replication^10^. *In vivo* DNA data writing therefore offers a route towards self-renewing and biologically maintained molecular memory.

Here, we describe <u>CON</u>jugative base-editing for in vivo <u>CO</u>ding and <u>R</u>ecording of <u>D</u>ata (CONCORD), a bacterial DNA data writing platform. A recipient “Writer” strain expresses the base editor and carries a designated writing sequence, while reusable donor “Guide” strains each carry a conjugative sgRNA plasmid that addresses a target in the Writer cell. This setup decouples the writing machinery from the sgRNA signal and enables programmable multiplexed writing of binary information directly into the bacterial genome by combining selected Guides with a Writer in a mixed-strain population. Using this framework, we encoded more than 45 kilobits of information, demonstrate orthogonal writing chemistries, pooled storage, random access and high-fidelity data recovery. CONCORD achieves scalable *in vivo* DNA writing through modularly delivered information and multiplexed data writing. We establish conjugation-coupled base editing as a programmable strategy for heritable information writing in bacteria and illustrate its use for sustainable DNA data storage.

## Results

### Conjugative delivery of sgRNAs enables programmable binary writing in bacterial genomes

To establish programmable *in vivo* DNA writing in *E. coli*, we combined CRISPR-guided base editing with bacterial conjugation. Base editors (BEs) are fusion proteins comprising a catalytically impaired CRISPR-Cas9 nickase (nCas9) and a nucleotide deaminase, enabling targeted and efficient base conversions without double-strand breaks **(Fig. 1a)**. A single guide RNA (sgRNA) with a short 20-nt spacer is the encoding signal used by nCas9 to guide the deaminase to a corresponding homologous target sequence on the genome. In our binary information encoding scheme, the unedited genomic base is assigned the state “0”, whereas the programmed base conversion is assigned the state “1”. By directing base editing to specific target positions, binary information is written directly into genomic DNA.

**Figure 1:**
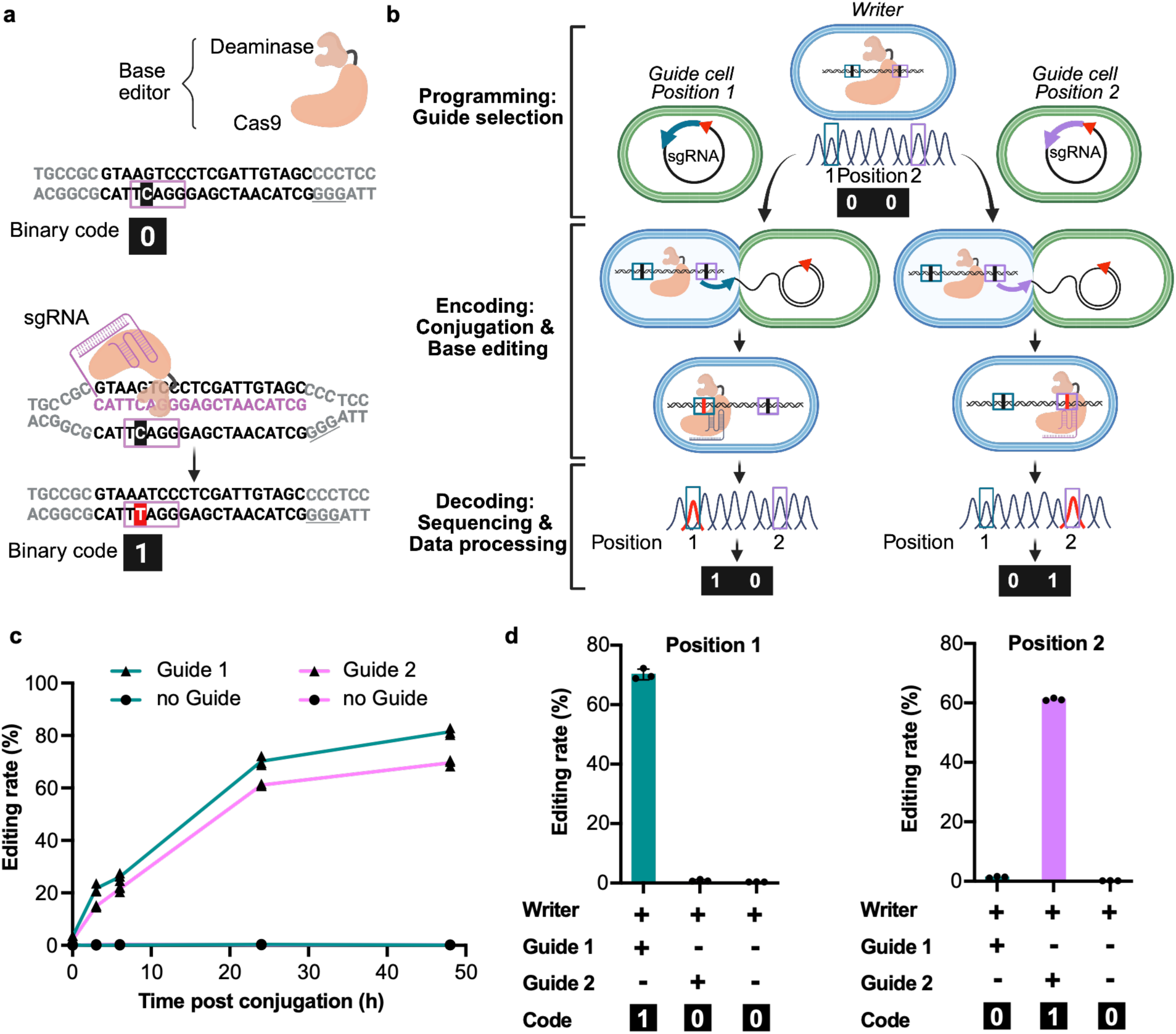
Programmed base editing at defined genomic target positions writes and stores binary data in DNA in vivo. **(a)** Binary encoding principle using base editing. A CBE converts a target cytosine (highlighted in black) within the deaminase target window (purple frame) to a thymine (highlighted in red) in presence of a sgRNA. The programmed base conversion is defined as a “1”, while an unmodified base is defined as “0”. The protospacer adjacent motif (PAM) sequence required by nCas9 is underlined. **(b)** Programmed binary encoding, addressing two positions in *lacZ* in the Writer genome. Writer cells express the base editor, while the conjugation-competent Guide cells (F+) each carry one sgRNA plasmid with an origin of transfer (red triangle). Horizontal transfer delivers the sgRNA to recipient Writer cells (F–), directing edits at target positions 1 or 2, respectively. **(c)** Timeline of the writing process. Editing rates were recorded at position 1 (green) and position 2 (purple) following conjugation with the corresponding Guides or no Guide. Encoding reactions were performed in biological triplicates and means are shown. **(d)** Editing rates of independent programmed reactions at positions 1 and 2 24 hours post conjugation were used to decode the written DNA sequence back to binary code. Encoding reactions were performed in biological triplicates and means with standard deviation are shown.

To achieve programmable encoding, we separate the writing machinery from the addressing signal across two bacterial cell types and utilize bacterial conjugation to control the information writing. A recipient ‘Writer’ strain expresses the base editor, while donor ‘Guide’ strains each carry a plasmid that contains an origin of transfer as well as single sgRNA targeting a specific position within the Writer genome **(Fig. 1b)**. Guide cells are conjugation-competent F⁺ bacteria equipped with an engineered non-transferrable IncP plasmid. This enables efficient horizontal gene transfer of the sgRNA plasmid to the recipient F⁻ Writer cell via bacterial mating without transfer of the conjugative machinery^32,33^. In this setup, the selection of the Guide strain for mating with the Writer determines which genomic position is addressed in the Writer genome. Following transfer, sgRNA transcription in the Writer directs site-specific editing, thereby converting DNA transfer into programmable genomic writing **(Fig. 1c)**. During post-conjugation recovery, we select for Writers that acquired the sgRNA plasmid.

We first tested this principle using a cytosine base editor and two sgRNAs targeting distinct positions within the non-essential lacZ locus. After writing, the target loci were sequenced and editing rates were quantified as the fraction of sequencing reads carrying the programmed C→T substitution at target bases. The deaminase-mediated base substitution depends on DNA repair. Consequently, editing rates increased over time in the cell growth recovery period, exceeding 60% for the respective positions within 24 hours. Position 1 was edited only in the presence of Guide 1, and position 2 only in the presence of Guide 2, whereas no edits were detected in the absence of Guide strains **(Fig. 1d)**. Thus, selective conjugative delivery of guide information enabled programmable writing of defined binary states in the bacterial genome.

### Increasing data density and improving the signal-to-noise ratio

Having established single-position encoding, we next sought to increase information density by combining multiple Guide cells with the Writer in a pooled conjugation strain mixture **(Fig. 2a)**. We identified six sgRNAs with high predicted editing efficiency scores within the *lacZ* locus, allowing up to 6 bits to be written in parallel **(Supplementary Data 1)**. Moreover, we introduced a PCR barcoding step to amplify and label *lacZ* loci from each mixture after writing for pooled information storage and sequencing. To recover the encoded information, we aggregated per-site edits across all sequencing reads from one mixture into a single binary code output reflecting all parallel encoding events. With this two-dimensional scaling strategy, we encoded 144 bits across the six positions across 24 conjugation populations **(Fig. 2b-c, Extended Data Figure 1)**.

**Figure 2:**
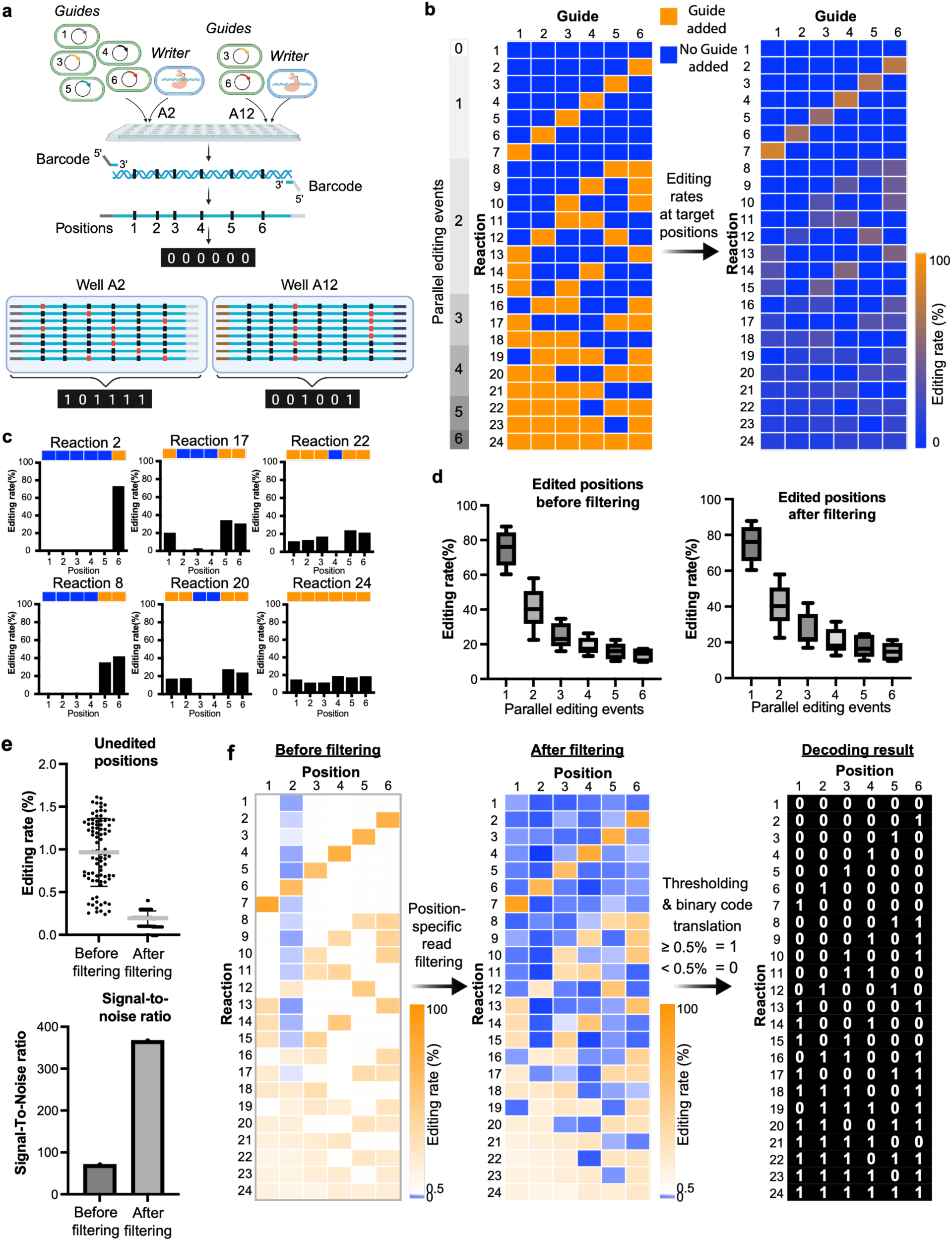
Scaling data density and reducing noise. **(a)** Multiple Guide strains are combined with a Writer in one population. A dual-index PCR barcode labels each population for pooled storage as barcoded amplicons. 6-bit codes were encoded in a 96-well plate. **(b)** Twenty-four 6-bit codes with up to six Guides per population encoded a total of 144 bits. For each population (left) editing rates were calculated at six *lacZ* target positions (right), corresponding to Guides 1-6. Each encoding reaction was performed once and editing rates recorded after 24h are shown. **(c)** Examples of programmed patterns and editing rates for 6 of 24 populations (full set in Extended Data Figure 1). **(d)** Editing rates at programmed “1”s across the 24 populations grouped by number of parallel editing events before and after position-specific read filtering. Standard box plots with median marked. **(e)** Distributions of apparent editing at programmed “0”s and signal-to-noise ratio across the 24 populations before and after position-specific filtering. Individual values, mean and standard deviation are shown for unedited position. Signal-to-noise ratio was calculated as: (‘mean of programmed 1s’ – ‘mean of programmed 0s’)/’standard deviation of programmed 0s’, resulting in a single signal-to-noise value before and after filtering. **(f)** Effect of position-specific read filtering at all 24 populations and decoding results using a 0.5% editing rate threshold.

Because recipient Writers remain F− due to the non-transferrable F-plasmid, Writers can experience repeated sgRNA deliveries. About 11% of reads were edited at two distinct target sites, indicating only a small fraction of simultaneous or consecutive conjugal deliveries from distinct Guide cells to the same Writer cell within the 1-hour conjugation time frame (**Extended Data Figure 2**). This shows that multiplexed writing within the same cell is feasible. Instead of multiplexing writing across genomic positions within the Writer cells multiplexing across the entire Writer population offers a more straight-forward path to data density increase. A population-based multiplexed writing strategy needs to take into account that the per-position editing rates decreases with increasing number of Guide strains, consistent with competition among donor strains for transfer into recipient cells **(Fig. 2d)**. We observed a mean apparent C→T base substitution of 1.0±0.4% at unedited positions (programmed “0”s), which we denominate as noise “editing rate” due to the intrinsic nanopore sequencing error^34,35^ **(Fig. 2e)**. To increase the number of parallel edits in a Writer population with distinct Guide cells and thus the number of bits encoded in the Writer population, we need to ensure that the editing rate signal remains above the sequencing noise. We developed a position-specific read filtering process that evaluates per-base sequencing quality independently at each target position for each read. A read may be discarded for editing rate determination at one position but included in the editing rate determined at another position due to varying quality across the read. This filtering approach reduced noise editing rates to 0.2±0.1% and significantly improved the signal-to-noise ratio while retaining 90% of the read depth **(Fig. 2d-e)**. Selectively excluding low-confidence base calls at target positions had a strong effect on programmed “0”s but minimally affected the signal editing rates at programmed “1”s. Following this filtering process we applied a 0.5% editing rate threshold to distinguish programmed binary states and correctly decoded all 144 bits **(Fig. 2f)**.

### Orthogonal base editors and synthetic writing sequences expand the encoding repertoire and writable sequence space

To expand the addressable sequence space for data writing, we systematically evaluated eight base editor variants with distinct protospacer adjacent motif (PAM) and deaminase specificities **(Fig. 3a, Supplementary Data 9-16)**. Editors that recognize different PAMs allow using one sgRNA to address duplicated protospacer sequences placed next to distinct PAMs, while editors with orthogonal deaminases encode distinct base substitution at each target site with the same sgRNA. Thus, base editors with orthogonal PAM specificities can expand the number of target positions addressable with the same set of Guide cells for increased data density, while base editors with distinct encoding patterns can for example be utilized to improve data recovery through redundancy encoding.

**Figure 3:**
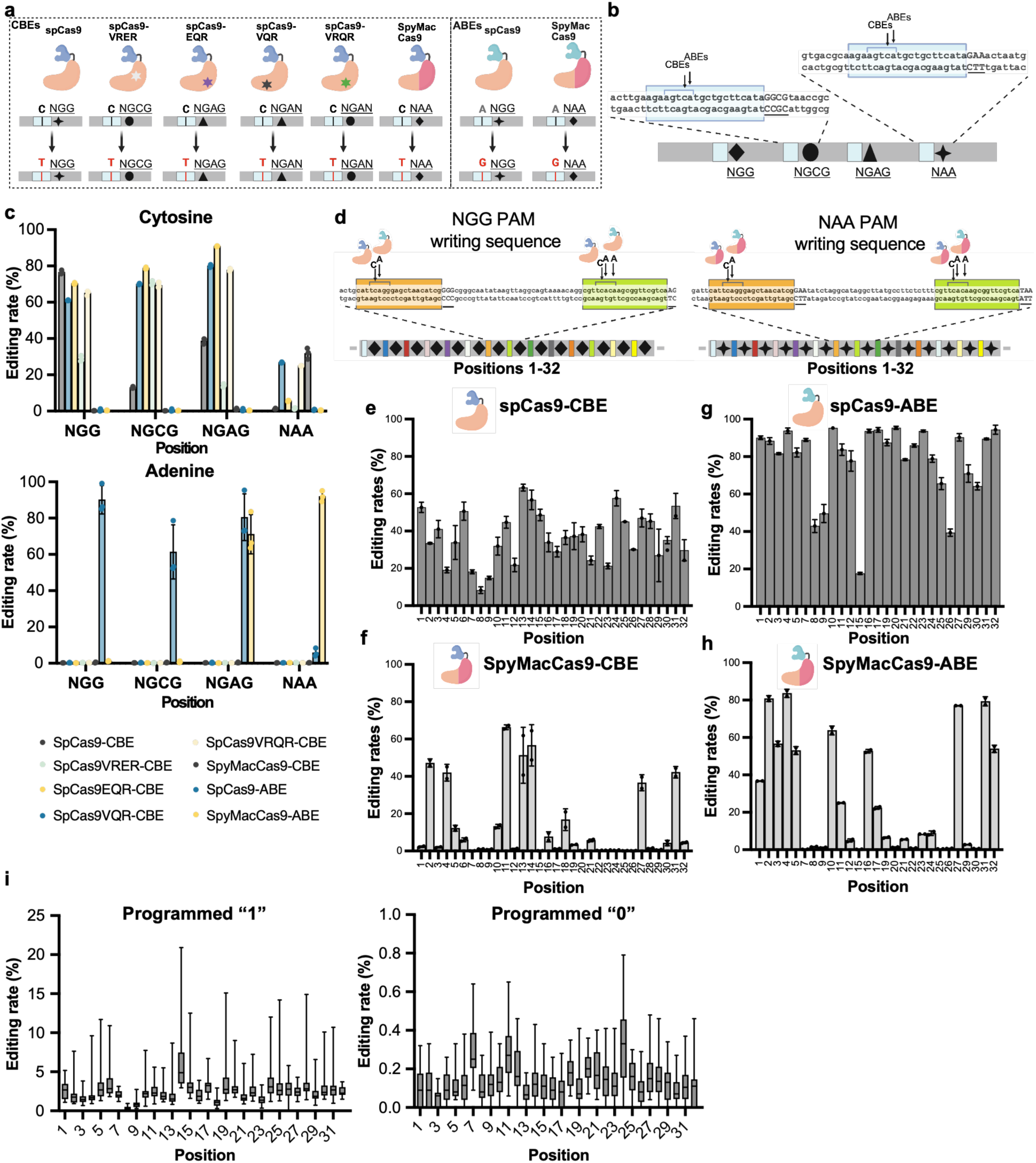
Orthogonal editors and compact synthetic writing sequences expand the encoding space. **(a)** Eight base editors screened for orthogonality, showing main PAM specificity and base conversion^38,39^. SpCas9 (wild type), SpCas9 mutants, and SpyMacCas9 were combined with APOBEC (dark blue) or TadA-8e (light blue). **(b)** Test cassette with four repeats of a protospacer (blue box), each followed by a distinct PAM (underlined). The deaminase main targeting window (bases 4-8) and target bases are indicated. **(c)** Editing rates at the four positions 24 hours after sgRNA delivery quantified for each Writer. Each experiment was performed in biological triplicates, means and standard deviation are shown. **(d)** Synthetic writing cassettes with 32 protospacers (rectangles), flanked by either NGG (diamond) or NAA (star) PAMs. Editing rates per position for **(e)** SpCas9-CBE and **(g)** SpCas9-ABE recorded on the NGG-cassette and for **(f)** SpyMacCas9-CBE and **(h)** SpyMacCas9-ABE on the NAA-cassette. Each experiment was performed in biological duplicates, means and standard deviation are shown. **(I)** 32-bit codes were encoded with SpCas9-CBE across 48 populations. Editing rates at programmed “1” and “0” states, editing rates for each grouped by position. Standard box plots with median marked.

We compared eight base editor variants: six cytosine base editors (CBEs) with an APOBEC deaminase and two adenine base editors (ABEs) with a TadA-8e deaminase. The deaminases were coupled to nCas9s with a range of PAM specificities: wild-type SpCas9, one of four SpCas9 mutants or the fusion SpyMacCas9. To directly compare editing efficiencies and PAM specificities we designed and genomically integrated a synthetic sequence with four repeats of a protospacer, each flanked by a distinct PAM, creating four target positions addressable with the same sgRNA **(Fig. 3b, Supplementary Data 17)**. After conjugative sgRNA delivery, we quantified C→T and A→G conversions at each position. Across this panel, ABEs showed higher editing efficiencies than their CBE counterparts, on average 2.7-fold higher for SpCas9-based editors and 6.7-fold higher for SpyMacCas9-based editors, but also a reduced PAM specificity, in line with previous studies in mammalian cells^36,37^ **(Fig. 3c)**. SpyMacCas9-ABE showed activity at the NGAG site, and SpCas9-ABE showed low activity at the NAA site, while their CBE counterparts did not edit those positions. SpCas9 PAM-relaxed mutants demonstrated decent base editing efficiencies but lower PAM specificity than reported in mammalian cells^38^. Overall, wild type SpCas9 and SpyMacCas9 showed most divergent PAM specificity within the tested set of PAM target sites. As expected, within the target positions, we observed no cross-base conversions, i.e. ABEs did not edit cytosines and CBEs did not edit adenines. To our knowledge, this provides the first direct comparison of several of these editors’ efficiency and specificity in *E. coli*.

We then designed a compact synthetic writing sequence containing 32 protospacers, condensing the writing space for 32 bits to approximately 1.7 kb **(Fig. 3d, Extended Data Figure 3, Supplementary Data 1, 18-19)**. Two versions of the writing sequence were synthesized and genomically integrated with identical protospacers but flanked either by an NGG or NAA PAM. Because the protospacers were identical, the same set of sgRNAs could be reused across positions, while the PAM context determined which editor could access the site. We measured editing efficiency at each position for SpCas9-CBE and SpCas9-ABE on the NGG cassette, and for SpyMacCas9-CBE and SpyMacCas9-ABE on the NAA cassette **(Fig. 3e-h, Extended Data Figure 4)**. We selected these four BEs due to their distinct and orthogonal base and PAM specificity. SpCas9-CBE and SpCas9-ABE showed broad activity across the NGG cassette, whereas the SpyMacCas9 editors were inactive at approximately half of the target positions and therefore not pursued further for large-scale encoding. Despite comparable predicted editing efficiencies, editing rates varied by position, with SpCas9-CBE typically yielding lower and more variable conversion than SpCas9-ABE. For each sgRNA, we selected the editable base with the highest efficiency as the canonical reporting position assessed for editing in subsequent analyses **(Supplementary Data 1)**. We did not observe off-target editing across the target positions for any sgRNA tested **(Extended Data Figure 4)**. While the specific editing window is editor- and nuclease-dependent, we observed a wider editing window for ABEs compared to their CBE counterparts **(Extended Data Figure 5)**.

Finally, to evaluate the practical upper limit of multiplexed writing we programmed 48 32-bit codes with SpCas9-CBE utilizing all 32 protospacers within the synthetic writing sequence. We confirmed that all positions except one were decodable with up to 32 parallel editing events using a 0.5% threshold after applying our position-specific filtering strategy **(Fig. 3i, Extended Data Figure 6)**. We later identified an 8-bp insertion at the origin of transfer of two sgRNA plasmids corresponding to the positions with especially low editing rates (positions 8 & 9), indicating a reduced conjugative sgRNA delivery rate rather than protospacer inaccessibility as the reason for the low observed editing efficiencies. We discarded Guide 8 for further encoding with SpCas9-CBE but kept both Guides for SpCas9-ABE encoding, because the higher ABE editing efficiency could compensate for the reduced delivery efficiency.

### Training and refining decoding of multiplexed base-editing-encoded data

Having established multiplexed writing, suitable target positions and orthogonal editors we next evaluated end-to-end encoding and decoding using a test dataset comprising two texts and a 20×8 pixel image **(Fig. 4a, Extended Data Figure 7)**. We converted the data to binary code and encoded it in biological triplicate using either the SpCas9-CBE Writer or the SpCas9-ABE Writer, with 31 and 27 Guides, respectively **(Supplementary Data 3)**. In total, 2,160 bits were encoded with each Writer, after which all writing sequenced were amplified, barcoded, pooled and sequenced.

**Figure 4:**
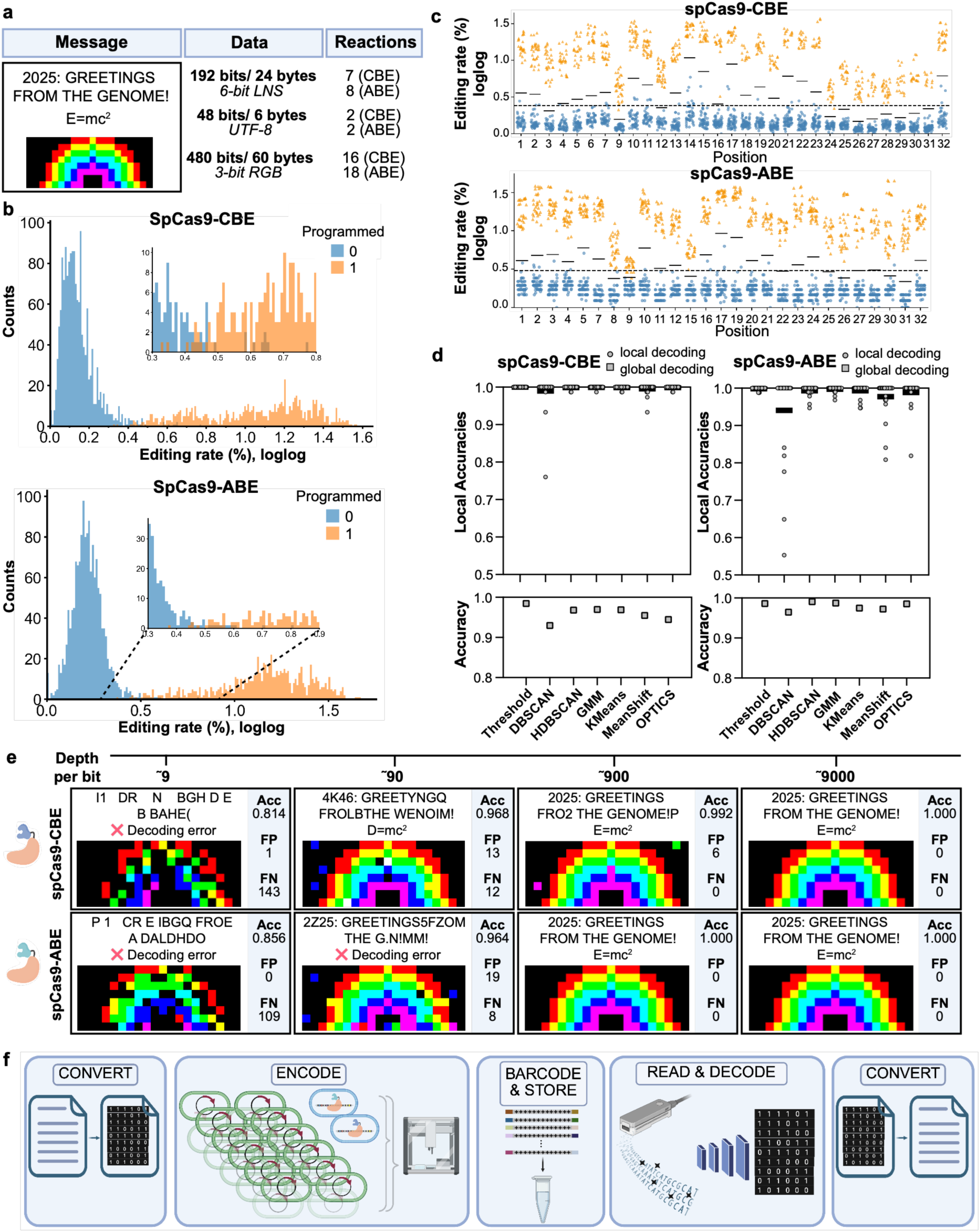
Testing and validating decoding strategies. **(a)** A test dataset was encoded in biological triplicates with SpCas9-CBE (31 bits per population) and SpCas9-ABE (27 bits per population). **(b)** Distribution of shifted double-log transformed editing rates for both Writers, labelled by their programmed binary state. Test datasets were encoded in biological triplicates, and all replicates’ editing rates are shown together in the plots. **(c)** Thresholds learned globally (dashed) and per position (solid black) for test datasets. Positions refer to the 32 sites on the NGG cassette. **(d)** Accuracies of data decoding by supervised learning (thresholds) as well as unsupervised learning (clustering algorithms: DBSCAN, HDBSCAN, GMM, KMeans, MeanShift, and OPTICS) approaches. Accuracy refers to the fraction of correctly decoded bits. Datasets were decoded by position (local) or as one dataset (global) and mean accuracy across all positions is depicted as black bars for local clustering. **(e)** Decoding results for biological triplicates of encoded data using trained local thresholds, with accuracies (Acc), false positives (FP, bits falsely identified as “1”) and false negatives (FN, bits falsely identified as “0”). **(f)** CONCORD end-to-end workflow, for in vivo writing, pooled storage, and decoding.

Editing rate distributions for programmed "1"s and "0"s were well separated overall, although some overlap remained **(Fig. 4b)**. We log-transformed editing rates and applied both supervised threshold-based methods and unsupervised clustering approaches to facilitate decoding of this larger dataset. Because editing efficiencies differed by target position and by editor, position-specific thresholds achieved better decoding results than a single global threshold **(Fig. 4c-d, Supplementary Data 4 & 5)**. Among the unsupervised clustering algorithms tested by position (local clustering) or across all positions (global clustering), local Gaussian mixture modelling (GMM), HDBSCAN and K-means performed best. At sequencing depths of approximately 1,000 reads per template (containing 31 or 27 bits depending on the editor used) we could fully recover all encoded information across the triplicates for both Writers **(Fig. 4e, Supplementary Data 5)**. These experiments established that multiplexed *in vivo* DNA writing of up to 31 bits per population and pooled storage of over 2 kbits can be decoded reliably from nanopore sequencing data.

Importantly, in the process of encoding the test datasets, we automated most experimental steps on liquid handling robots and adapted the protocol to a 96-well format using microlitre-scale cellular input materials, which enabled parallelized encoding. We refer to the resulting workflow as CONCORD (<u>CON</u>jugative base-editing for in vivo <u>CO</u>ding and <u>R</u>ecording of <u>D</u>ata). CONCORD integrates *in vivo* binary data encoding, barcoding, pooled storage, and decoding into a high-throughput, scalable end-to-end workflow that requires only a Writer strain and a reusable set of Guide strains as biological inputs **(Fig. 4f, Extended Data Figure 8** & **9)**.

### Blind decoding, redundancy encoding, and random-access support large-scale information recovery

To demonstrate the capacity of CONCORD for cold data storage, we encoded two relevant archival objects: a 1,134-character greeting from NASA’s Voyager Golden Record (Voyager), and a photograph of the Navajo petroglyphs from Crow Canyon converted to a 97×66-pixel grayscale image including a short text with image specifications (Petroglyph) **(Fig. 5a)**. The Voyager dataset was encoded independently with both SpCas9-CBE and SpCas9-ABE, creating distinct encoding patterns due to their orthogonal editing chemistries. The Petroglyph dataset was only encoded with SpCas9-CBE. Across these experiments, 39,348 bits were written.

**Figure 5:**
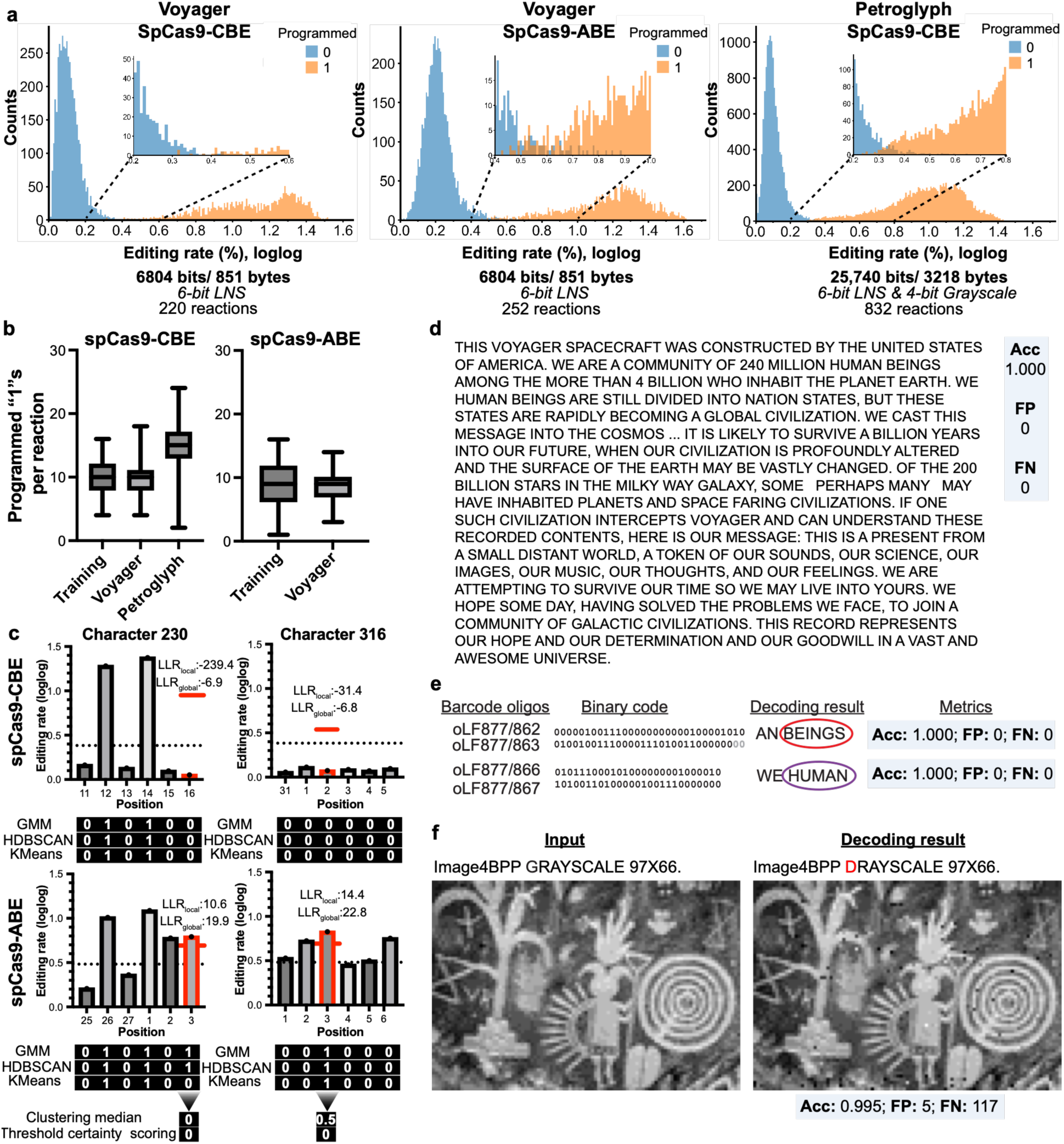
Large-scale CONCORD encoding of significant artifacts with blind decoding, redundancy, and random access. **(a)** Distributions of shifted double-log-transformed editing rates for two large datasets: A 1,134-character text (Voyager) encoded twice with two different Writers and a 97×66-pixel image plus 22-character text encoded once (Petroglyph). **(b)** Comparison of the number of programmed “1”s per population in large-scale datasets versus the test dataset from Figure 4. Box plots of all programmed “1”s editing rates with median marked. **(c)** Strategy for solving discrepancies in decoding of redundancy encoded data. The median over 6 local clustering predictions from GMM, HDBSCAN, and KMeans was formed for all bits of uncertain characters and certainty scores (LLR) were computed based on distance of editing rates from global or local thresholds at the respective positions. In case of uncertain clustering (median of 0.5), a threshold-based certainty scoring approach was followed taking all four certainty scores into account for an uncertain bit. Plots show single value editing rates since all encodings were performed in single biological replicates. **(d)** Voyager greeting recovered from redundancy encoding with SpCas9-CBE and SpCas9-ABE. **(e)** Random access recovery of selected words from pooled Voyager datasets (red circle, SpCas9-CBE; purple circle, SpCas9-ABE) using barcode-specific primers. **(f)** Petroglyph image plus introductory text: Input and recovered output.

We found decoding by local clustering the best approach for blind decoding of these larger datasets, outperforming threshold-based decoding in accuracy **(Extended Data Figure 10, Supplementary Data 7)**. Previously trained thresholds could not adapt to changes in dataset parameters that affect the overall editing rates, in particular the proportion of programmed “1”s per population. This was most apparent in the Petroglyph dataset, which has a higher proportion of “1”s (49%) compared to the test dataset used to train the thresholds (32%) **(Fig. 5b, Supplementary Data 3 & 6).** Rather than relying on fixed thresholds for decoding clustering methods adapt to new datasets, and local clustering overall achieved best decoding accuracies. For optimal decoding robustness, we used a consensus strategy based on majority voting across the three best-performing clustering algorithms: GMM, HDBSCAN, and K-means.

Encoding the Voyager dataset twice with two orthogonal Writers allowed us to cross-validate ambiguous data. Four of the 1,134 characters differed in the redundantly encoded dataset after conversion of binary output to characters. We resolved these discrepancies in a 2-step evaluation of the underlying bits, by i) combining the six local clustering predictions and ii) computing threshold-based certainty scores. For example, clustering did not provide a conclusive decision for character 316, but the threshold-based evaluation evaluated the SpCas9-CBE datapoint with a higher certainty **(Fig. 5c, Extended Data Figure 11)**. This approach yielded complete recovery of the Voyager greeting and demonstrated that orthogonal writing chemistries can provide robustness for data recovery through redundancy encoding **(Fig. 5d)**.

We next asked whether individual records could be retrieved selectively from pooled amplicon archives. Using primers complementary to population-specific barcodes we amplified seven selected words from the pooled Voyager DNA datasets **(Fig. 5e, Extended Data Figure 12, Supplementary Data 6)**. The correct recovery of all seven words demonstrates successful random-access retrieval from pooled data archives, an important feature of practical data storage.

Finally, we decoded the Petroglyph data by majority-vote local clustering **(Fig. 5f, Supplementary Data 7)**. The image and accompanying text were recovered at 99.5% accuracy. The high data recovery rate of this dataset despite a lack of replicate or redundancy encoding and without relying on computational error correction shows the high encoding and decoding fidelity of CONCORD. The image was clearly recognizable with all details and only few remaining errors affecting 118 pixels of the 6402-pixel image. In summary, these experiments demonstrate that our conjugation-coupled bacterial base editing system can be used for high-fidelity information writing and recovery in bacteria. Compared to other synthesis-free DNA data storage methods, CONCORD excels in writing fidelity and scalability **(Supplementary Data 20)**.

## Discussion

In this study, we developed a bacterial platform that couples conjugative sgRNA delivery to CRISPR base editing for programmable *in vivo* DNA writing. By separating the editing machinery from the guide signal across recipient and donor strains, respectively, CONCORD converts a native bacterial DNA-transfer process into an addressable writing system. Information is encoded by selecting combinations of bacterial Guide strains that direct heritable nucleotide changes at defined genomic positions in a Writer strain and not by chemically synthesizing new DNA molecules. To facilitate information recovery and expand the writable space we genomically integrated a compact sequence with 32 protospacers in the Writer. A central feature of CONCORD is that the writing process is distributed across interacting bacterial populations. The Writer strain supplies the genomic substrate and editing machinery, whereas Guide strains specify which positions are edited through conjugative transfer. This separation has interesting practical and conceptual features. Practically, Guide strains can be reused and combined in different combinations to generate distinct binary code outputs without rebuilding the Writer. Conceptually, conjugation is repurposed as a programmable process for directed and heritable genome modifications. Conjugative DNA transfer is used to control which genomic states are installed in recipient cells. This setup establishes a modular framework for information writing in bacteria.

We found that we could increase information density through multiplexed writing, combining multiple Guide cells with the Writer in a mixed population. While the delivery of multiple sgRNAs to the same cell only occurred in a fraction of Writer cells, we could capture multiplexed writing by parallel conjugation events across the Writer cell population. We have developed a sequencing read denoising algorithm that incorporates a base-specific quality filtering step before determining the editing rates to improve the signal-to-noise ratio, addressing the decrease in editing signal with increasing number of Guides. We also found unsupervised clustering by position the best approach for separating programmed “1”s from programmed “0”s, reflecting the variation in editing efficiencies across target positions. Conjugation delivery efficiency also affected editing efficiency, as sgRNA plasmid with impaired origin of transfer correlated with significantly decreased editing rates. Thus, the capacity of parallel writing is shaped not only by editor activity, but also by the population dynamics of conjugative delivery.

Next, we identified base editors with orthogonal encoding repertoires. SpCas9-CBE and SpCas9-ABE showed the highest activity across all 32 protospacers flanked by NGG PAMs on a synthetic writing cassette. Notably, protospacers were selected based on predicted editing efficiencies for SpCas9-CBE and not all of them contained an adenine within the editing window, which reduced the Guide set usable with ABE. Moreover, while SpyMacCas9-based editors showed very good PAM orthogonality with SpCas9-based editors they only showed editing activity at about half of the protospacers on an analogous NAA PAM writing cassette. We conclude that editor-specific target activity, editing windows and the presence of targetable bases are important aspects to consider when using multiple orthogonal editors for DNA writing. Here, we use editors with orthogonal deaminase activities for redundancy encoding, improving data recovery through data validation across two distinct encoding patterns. One could also imagine expressing orthogonal editors simultaneously or sequentially in the same cell using the same set of Guides on an expanded writing sequence for further increased information density and reduced sequencing coverage requirements.

In summary, we have demonstrated that bacterial conjugation and base editing can be combined into a robust, end-to-end system for genomic information writing and recovery. We encoded more than 45 kilobits of binary data into bacterial DNA, achieved complete recovery of redundantly or repeatedly encoded information and near-complete recovery of a single-encoded image, and demonstrated random-access retrieval from pooled archives. We implemented multiplexed *in vivo* DNA data encoding with automation integration and demonstrated CONCORD’s potential for high-throughput information encoding. Thereby we overcome key challenges of DNA data encoding: the reliance on chemically synthesized oligonucleotides and purified enzymes of *in vitro* strategies, as well as the inefficiency and lack of scalability of *in vivo* strategies^10^. While we apply CONCORD here for DNA data storage, the system provides a means to program stable genomic memory through controlled interactions between bacteria in a mixed population of bacterial strains. This may be useful wherever defined cell-cell encounters are coupled to persistent genetic outcomes, including biological recording, engineered memory and cellular computing. To this end, further extension of the platform beyond *E. coli*, as well as probing long-term evolutionary stability of encoded states would be relevant next steps.

## Acknowledgements

We thank W.Z. Teo and C.P. Han and the National University of Singapore Biofoundry for help and access to automation platforms, and J. Fredens for helpful feedback on the manuscript.

## Funding

This work was supported by the Lee Kuan Yew Postdoctoral Research fellowship awarded to L.F.H.F. by the National University of Singapore, AcRF Tier 2 by the Ministry of Education (T2EP30221-0035 to C.L.P), Singapore and the National Centre for Engineering Biology (NRF-MSG-2023-0003 to C.L.P) by The National Research Foundation, Singapore.

## Author contributions

C.L.P supervised and administered the project. L.F.H.F. designed the project and performed all CONCORD experiments. L.F.H.F. and Z.L. developed the denoising strategy and algorithm. Z.L. developed barcoding strategy and supported Nanopore sequencing. W.L. developed the Bitconverter and supported decoding strategies. Y.L. supported threshold training. L.F.H.F wrote the original manuscript and all authors reviewed and edited the manuscript. ICMJE guidelines were followed.

## Competing interests

The authors declare no competing interests. The National University of Singapore has filed a patent application covering CONCORD.

## Supplementary

Materials and Methods

Extended Data Figures 1-11

Supplementary Data 1-21

## Notes

### Competing Interest Statement

The authors have declared no competing interest.

## References

1 Sanchez-Romero, M. A. & Casadesus, J. The bacterial epigenome. Nat Rev Microbiol 18, 7–20 (2020). 10.1038/s41579-019-0286-2

2 Scanlon, K., Shanahan, F., Ross, R. P. & Hill, C. Exploring the concept of bacterial memory. Nat Microbiol 10, 3049–3058 (2025). 10.1038/s41564-025-02185-3

3 Arnold, B. J., Huang, I. T. & Hanage, W. P. Horizontal gene transfer and adaptive evolution in bacteria. Nat Rev Microbiol 20, 206–218 (2022). 10.1038/s41579-021-00650-4

4 Barrangou, R. et al. CRISPR provides acquired resistance against viruses in prokaryotes. Science 315, 1709–1712 (2007). 10.1126/science.1138140

5 Lim, C. K., Yeoh, J. W., Kunartama, A. A., Yew, W. S. & Poh, C. L. A biological camera that captures and stores images directly into DNA. Nat Commun 14, 3921 (2023). 10.1038/s41467-023-38876-w

6 Farzadfard, F. & Lu, T. K. Synthetic biology. Genomically encoded analog memory with precise in vivo DNA writing in living cell populations. Science 346, 1256272 (2014). 10.1126/science.1256272

7 Yim, S. S. et al. Robust direct digital-to-biological data storage in living cells. Nat Chem Biol 17, 246–253 (2021). 10.1038/s41589-020-00711-4

8 Loveless, T. B. et al. Open-ended molecular recording of sequential cellular events into DNA. Nat Chem Biol 21, 512–521 (2025). 10.1038/s41589-024-01764-5

9 Farzadfard, F. et al. Single-Nucleotide-Resolution Computing and Memory in Living Cells. Mol Cell 75, 769–780 e764 (2019). 10.1016/j.molcel.2019.07.011

10 Wang, S., Mao, X., Wang, F., Zuo, X. & Fan, C. Data Storage Using DNA. Adv Mater 36, e2307499 (2024). 10.1002/adma.202307499

11 Sadremomtaz, A., et al. Digital data storage on DNA tape using CRISPR base editors. Nat Commun 14, 6472 (2023). 10.1038/s41467-023-42223-4

12 Cabezon, E., Ripoll-Rozada, J., Pena, A., de la Cruz, F. & Arechaga, I. Towards an integrated model of bacterial conjugation. FEMS Microbiol Rev 39, 81–95 (2015). 10.1111/1574-6976.12085

13 Lederberg, J. & Tatum, E. L. Gene recombination in Escherichia coli. Nature 158, 558 (1946). 10.1038/158558a0

14 Anzel, A., Heider, D. & Hattab, G. The visual story of data storage: From storage properties to user interfaces. Comput Struct Biotechnol J 19, 4904–4918 (2021). 10.1016/j.csbj.2021.08.031

15 Appuswamy, R. in Encyclopedia of Big Data Technologies (eds Sherif Sakr & Albert Y. Zomaya) 435–443 (Springer International Publishing, 2019).

16 Erlich, Y. & Zielinski, D. DNA Fountain enables a robust and efficient storage architecture. Science 355, 950–954 (2017). 10.1126/science.aaj2038

17 Zhirnov, V., Zadegan, R. M., Sandhu, G. S., Church, G. M. & Hughes, W. L. Nucleic acid memory. Nat Mater 15, 366–370 (2016). 10.1038/nmat4594

18 Organick, L. et al. Probing the physical limits of reliable DNA data retrieval. Nat Commun 11, 616 (2020). 10.1038/s41467-020-14319-8

19 Kjaer, K. H. et al. A 2-million-year-old ecosystem in Greenland uncovered by environmental DNA. Nature 612, 283–291 (2022). 10.1038/s41586-022-05453-y

20 Grass, R. N., Heckel, R., Puddu, M., Paunescu, D. & Stark, W. J. Robust chemical preservation of digital information on DNA in silica with error-correcting codes. Angew Chem Int Ed Engl 54, 2552–2555 (2015). 10.1002/anie.201411378

21 Lim, C. K., Nirantar, S., Yew, W. S. & Poh, C. L. Novel Modalities in DNA Data Storage. Trends Biotechnol 39, 990–1003 (2021). 10.1016/j.tibtech.2020.12.008

22 Church, G. M., Gao, Y. & Kosuri, S. Next-generation digital information storage in DNA. Science 337, 1628 (2012). 10.1126/science.1226355

23 Goldman, N. et al. Towards practical, high-capacity, low-maintenance information storage in synthesized DNA. Nature 494, 77–80 (2013). 10.1038/nature11875

24 Palluk, S. et al. De novo DNA synthesis using polymerase-nucleotide conjugates. Nat Biotechnol 36, 645–650 (2018). 10.1038/nbt.4173

25 Chen, Y. J. et al. Quantifying molecular bias in DNA data storage. Nat Commun 11, 3264 (2020). 10.1038/s41467-020-16958-3

26 Lee, H. et al. Photon-directed multiplexed enzymatic DNA synthesis for molecular digital data storage. Nat Commun 11, 5246 (2020). 10.1038/s41467-020-18681-5

27 Hao, M. et al. A mixed culture of bacterial cells enables an economic DNA storage on a large scale. Commun Biol 3, 416 (2020). 10.1038/s42003-020-01141-7

28 Shipman, S. L., Nivala, J., Macklis, J. D. & Church, G. M. CRISPR-Cas encoding of a digital movie into the genomes of a population of living bacteria. Nature 547, 345–349 (2017). 10.1038/nature23017

29 Hou, Z. et al. "Cell Disk" DNA Storage System Capable of Random Reading and Rewriting. Adv Sci (Weinh) 11, e2305921 (2024). 10.1002/advs.202305921

30 Zhang, C. et al. Parallel molecular data storage by printing epigenetic bits on DNA. Nature 634, 824–832 (2024). 10.1038/s41586-024-08040-5

31 Tabatabaei, S. K. et al. DNA punch cards for storing data on native DNA sequences via enzymatic nicking. Nat Commun 11, 1742 (2020). 10.1038/s41467-020-15588-z

32 Zurcher, J. F. et al. Continuous synthesis of E. coli genome sections and Mb-scale human DNA assembly. Nature 619, 555–562 (2023). 10.1038/s41586-023-06268-1

33 Fredens, J. et al. Total synthesis of Escherichia coli with a recoded genome. Nature 569, 514–518 (2019). 10.1038/s41586-019-1192-5

34 Sanderson, N. D. et al. Comparison of R9.4.1/Kit10 and R10/Kit12 Oxford Nanopore flowcells and chemistries in bacterial genome reconstruction. Microb Genom 9 (2023). 10.1099/mgen.0.000910

35 Luo, J. et al. Systematic benchmarking of nanopore Q20+ kit in SARS-CoV-2 whole genome sequencing. Front Microbiol 13, 973367 (2022). 10.3389/fmicb.2022.973367

36 Hu, J. H. et al. Evolved Cas9 variants with broad PAM compatibility and high DNA specificity. Nature 556, 57–63 (2018). 10.1038/nature26155

37 Kleinstiver, B. P. et al. High-fidelity CRISPR-Cas9 nucleases with no detectable genome-wide off-target effects. Nature 529, 490–495 (2016). 10.1038/nature16526

38 Kleinstiver, B. P. et al. Engineered CRISPR-Cas9 nucleases with altered PAM specificities. Nature 523, 481–485 (2015). 10.1038/nature14592

39 Chatterjee, P. et al. A Cas9 with PAM recognition for adenine dinucleotides. Nat Commun 11, 2474 (2020). 10.1038/s41467-020-16117-8

