## Supplementary Materials for "High-throughput *in vivo* DNA data encoding by multiplexed conjugation and orthogonal base editing"

### Materials & Methods

General methods.

All plasmids and oligonucleotides used in this study are listed in **Supplementary Data 2.** Plasmids were obtained from Addgene, or constructed in this study. All gBlocks were synthesized by Integrated DNA Technologies, and oligonucleotides were synthesized by Integrated DNA Technologies or Bio Basic Asia Pacific. *E. coli* was grown in LB medium (Bio Basic Asia Pacific) or on LB agar (Bio Basic Asia Pacific). Antibiotics used in this study for selection where appropriate are: kanamycin (50 µg/mL), ampicillin (100 µg/mL), streptomycin (200 µg/mL), apramycin (50 µg/mL), chloramphenicol (20 µg/mL), tetracycline (10 µg/mL), and hygromycin (200 µg/mL). Moreover we used 4-chlorophenylalanine (2.5 mM) for negative selection with *pheS*T251A_A294G (referred to as *pheS** throughout this study). Isopropylthio-β-galactoside (IPTG, 1 mM) or L-arabinose (0.5% w/v) were used to induce expression of proteins under T7 or araBAD promoter, respectively.

We prepared electrocompetent cells by growing up *E. coli* overnight at 37˚C under shaking in 10 ml LB with appropriate antibiotics if applicable. The dense overnight culture was inoculated 1:50 fold into 200 ml of the same medium. The culture was grown at 37˚C under shaking to $OD_{600}$ = 0.5, and then incubated on ice for 20 min. Cells were spun down at 4000 x g for 10 min at 4 °C. The supernatant was discarded, and the cell pellet was resuspended in 20 ml cold Milli-Q filtered water and then spun down. Cells were washed once more with cold Milli-Q filtered water followed by a wash with cold 16% (w/v) glycerol. Finally, the cell pellet was resuspended in 400 µl 16% glycerol and used immediately for transformation by electroporation, or frozen and stored at -80°C. We prepared chemically competent cells similarly, growing a 50 ml culture from a dense overnight culture at 37˚C to $OD_{600}$ = 0.5, then incubating the culture on ice for 20 min. Cells were spun down and the pellet resuspended in 20 ml cold 0.1 M $CaCl_2$ and incubated on ice for 30 min. After this incubation, cells were spun down, resuspended in 1.5 ml 16% glycerol and used immediately for heat shock transformation (1 min at 42˚C), or frozen and stored at -80°C.

Construction of plasmids

Cloning of all plasmids was carried out in *E. coli* DH10b cells. Most plasmids were constructed by Gibson assembly according to the manufacturer's protocol with NEBuilder HiFi DNA Assembly Master Mix (New England Biolabs). Fragments for the plasmid construction were amplified by PCR from template plasmids, gBlocks, or genomic DNA as specified in **Supplementary Data 2**, with 20 bp overlap regions between fragments. PCR reactions were performed using Q5® High-Fidelity Master-Mix (New England Biolabs). The Gibson assembly mixtures were transformed into chemically competent *E. coli* DH10b, and cells were plated on LB agar with antibiotics selecting for the constructed plasmid. Following overnight incubation we verified correct assembly by genotyping on 2-4 selected colonies by PCR with One*Taq*® Quick-Load® 2X Master Mix (New England Biolabs) and oligonucleotides flanking the overlap regions between fragments. For colonies that passed genotyping, we verified functionally relevant plasmid sections by Sanger sequencing performed by BioBasic or Axil Scientific. Sample preparation for PCRs and sequencing was done according to each companies' protocol.

We constructed the first sgRNA plasmid (pLF133b) from 4 fragments by Gibson assembly. All other sgRNA plasmids were constructed as 1-fragment assemblies from a fragment created by two oligonucleotides containing the new 20 bp spacer sequence homologous to the target protospacer in the oligonucleotide overlap region. Thus, all new sgRNA plasmid fragments were created by PCR with the same primer binding sites, enabling construction of novel sgRNA plasmids without further PCR parameter optimization. The plasmid map of sgRNA plasmid pFL208 is provided for reference in **Supplementary Data 8**.

For base editor plasmids we first constructed pLF177 (SpCas9-CBE) and pLF178 (Sp-Cas9-ABE) by 2-fragment Gibson assembly. pLF177 then served as template for the construction of SpCas9-mutant plasmids pLF180-pLF183, constructed as 2-fragment Gibson assemblies with the fragment introducing the point mutations amplified from a gBlock (gLF007-gLF010, respectively). For pLF184 (SpyMacCas9-CBE) and pLF210 (SpyMacCas9-ABE), we performed Golden Gate Assembly instead of Gibson assembly with the PaqCI restriction enzyme. The following reaction mixtures were prepared: ˜50 fmol vector PCR fragment, ˜100 fmol insert PCR fragment, 2 µl 10x T4 DNA ligase buffer (New England Biolabs), 0.8 µl PaqCI (New England Biolabs), 0.25 µl PaqCI activator (New England Biolabs), $H_20$ to 20 µl total volume. Golden Gate assembly reactions were performed on a thermocycler as follows: (37˚C, 1min -> 16˚C, 1min)x50 cycles, then 37˚C for 5min followed by 60˚C for 5 min. The entire reaction mixture was then transformed into chemically competent *E. coli* DH10b, and plated on agar with antibiotic selection. Plasmid verification was performed as described above, first by genotyping, then by sequencing. Final plasmids were verified by Nanopore sequencing performed in house using Flongle flow cells (Oxford Nanopore Technologies, ONT). Plasmid libraries were prepared with the Rapid Barcoding Kit 96 v14 (SQK-RBK114.96, ONT) as per protocol. Genbank files of all base editor plasmids are provided in **Supplementary Data 9-16.**

#### Selection of protospacers and design of synthetic writing sequences

We predicted protospacers with the 'Design CRISPR' tool on Benchling (Benchling) for base editing with SpCas9-CBE in *E. coli* MG1655. Protospacers were selected with high base editing scores (> 21) for C->T substitutions [1]. Editing predictions for other base editors used in this study were not available on Design CRISPR. Multiple predicted edits within the 20 bp sequence were accepted as long as the predicted base editing score for at least one edit was above 21. We selected six protospacers for the non-essential endogenous *lacZ* gene for initial CONCORD experiments editing the wild type *E. coli* genome directly. To expand and condense the writing space we then selected additional 27 protospacers predicted based on sequences from exogenous genes not present in the *E. coli* genome (human *alt1*, sfgfp, mcherry, proteus sp *gdh)*. All protospacers are listed in **Supplementary Data 1**.

We designed three synthetic sequences for writing and synthesized them as gBlocks: gLF3456 for orthogonality tests, gLF013 and gLF014 for data encoding. The synthetic sequence used for orthogonality screens of base editors was designed with four repeats of the protospacer AGAAGTCATGCTGCTTCATA, each flanked by a distinct PAM sequence (NGG, NGCG, NGAG, NAA). The synthetic writing sequences gLF013 and gLF014 were designed to contain 32 protospacers flanked by NGG PAM (gLF013) or NAA PAM (gLF014) and 30 bp spacing sequence in between protospacers selected from exogenous *alt1*, sfgfp, mcherry, *gdh* sequences. For each gBlock we first constructed a plasmid coupling the synthetic sequence to antibiotic selection markers to facilitate genomic integration by homologous recombination (pLF162b for gLF3456, pLF209 for gLF013, and pLF211 for gLF014). Genbank files of the synthetic writing sequences integrated at the *lacZ* locus are provided in **Supplementary Data 17-19**.

#### Construction of Writer and Guide strains

We constructed bacterial Writer strains by integrating synthetic writing sequences into the bacterial genome at the endogenous non-essential *lacZ* locus. All integrations were done by lambda-red mediated homologous recombination. To select for homogolous recombination we created a double-selection cassette of negative selection gene *rpsL* and positive selection gene *hygR* under the endogenous rpsL promoter synthesized as a gBlock (gLF001). Moreover, strains subjected to homologous recombination were transformed with pLF127, a plasmid carrying the lambda-red alpha/beta/gamma genes under araBAD promoter. To prepare cells for homologous recombination we prepared electrocompetent cells as described above, additionally inducing expression of the recombination machinery by adding L-arabinose at $OD_{600}$ = 0.2 for 1 hour during growth. Furthermore, we prepared linear DNA template for recombination by PCR amplification, typically with primers containing 50 bp homology to the target genomic integration sites. We electroporated ˜1 µg of purified PCR product into 50 µl of electrocompetent and induced cells harbouring pLF127. Cells were recovered for 3 hours in 50 ml LB medium and plated on LB agar with appropriate selection. Successful recombination was verified by colony PCR with oligonucleotides flanking the integration site and phenotypic assessment if applicable.

First, we created *E. coli* MG1655$^{rpsl\mathrm{K43R}}$, the base strain for all Writers, rendering *E coli* MG1655 streptomycin sensitive. We amplified *rpsL*K43R from the *E. coli* DH10b genome to create the template for recombination and plated cells on LB agar with streptomycin after recombination. Next, we integrated the T7 RNA polymerase under IPTG-inducible lacUV5 promoter in a two-step process (outlined in **Extended Data Figure 2**). First, we integrated the *rpsL-hygR* cassette amplified from gLF001 between endogenous *ybhC* and *ybhB* and selected with hygromycin for integration. Then we amplified the $P_{lac}$-*T7rnap* gene from the BL21(DE3) genome gene and selected with streptomycin in a second recombination experiment for scarless replacement of the *rpsL-hygR* cassette with the *T7rnap*. Integration was verified by genotyping and plaque formation with a test phage depending on T7 RNA polymerase activity (*λ*DE3 lysigenization kit, Novagen, 69734). The resulting strain is referred to as MG1655(T7RNAP). We transformed MG1655(T7RNAP) with pLF177 containing SpCas9-CBE for initial CONCORD experiments editing target positions in the native *lacZ* gene. We also subjected MG1655(T7RNAP) to further recombination, integrating synthetic writing sequences. To this end we created templates for recombination from pLF162b, pLF209 and pLF211 with primers containing 50 bp homology to the endogenous *lacZ* gene, the selected locus for integration. For integration of the gLF3456 synthetic sequence we selected with tetracycline for successful integration. For CONCROD experiments with 32 Guides we integrated gLF013 (NGG PAMs) coupled to *rpsL-hygR* and gLF014 (NAA PAMs) coupled to *pheS*-tetR*, selecting with hygromycin and tetracycline for integration, respectively. We chose double selection cassettes for the latter to enable simple further expansion of the writing sequence with negative selection [2]. The resulting strains were referred to as MG1655(T7RNAP)_RpLF162b, MG1655(T7RNAP)_RpLF209, and MG1655(T7RNAP)_RpLF211, and transformed with base editor plasmids.

For the construction of Guide cells, we transformed a non-transferrable F-plasmid with truncated origin of transfer (pJF146, kindly provided by Jason Chin)[2] into *E. coli* DH10b cells by electroporation. We then made the resulting cell chemically competent and introduced the sgRNA plasmid by heat shock transformation. Final Guide cells were selected for with apramycin (F-plasmid), and chloramphenicol (sgRNA plasmid). Guide cells are conjugation competent and able to transfer sgRNA plasmids into recipient cells due to an origin of transfer, but cannot transfer the F-plasmid.

<u>DNA data encoding with CONCORD</u>

To perform digital to biological data encoding with CONCORD, digital data was first converted to binary data using a character conversion standard. We used character encoding standards UTF-8, 3-bit RGB, and 4-bit grayscale, as well as a custom 6-bit character conversion termed 6-bit LNS (**Extended Data Figure 6**) in this study. The binary data was then converted to CONCORD populations for biological encoding by splitting the string into consecutive sections of for example 31 bits, depending on the number of bits encoded per cell. No padding was used except for completing the final section if necessary. The length of the binary string divided by the number of bits encoded per cell gives the number of populations required for DNA encoding. In this study we encoded 31 bits per cell with CBEs and 27 bits per cell with ABEs. For ABEs, editing rates were only recorded at 27 positions due to the lack of adenine presence at the remaining 5 positions **(Supplementary Data 1)**. We encoded across 96 populations per plate and used a fixed filling pattern across the plate (A1-A12, B1-B12, etc. for populations 1-12, 13-24, etc.). The Voyager text is encoded in 220 populations (2.3x 96-well plates) with SpCas9-CBE and 252 populations (2.6x 96-well plates) with SpCas9-ABE, and the Petroglyph dataset is encoded in 832 populations (2.6x 96-well plates).

We developed Bitconverter, a freely available browser-based web application implemented in Python using the Streamlit framework deployed on Hugging Face Spaces (https://huggingface.co/spaces/EngBioNUS/bitconverter). The encoding tool converts digital to binary data, and produces a binary matrix of the populations required for encoding grouped by (adjustable) target positions.

CONCORD data encoding utilizes Writer and Guide bacterial strains, prepared as described above. Guide cells are conjugation-competent donor cells equipped with a non-transferrable F-plasmid and an sgRNA plasmid with an *oriT* sequence. Writer cells are MG1655(T7RNAP) derivate cells equipped with a base editor plasmid and function as recipients during conjugation. Volumes required of each cellular culture depend on the number of encoding populations as well as the equipment used and need to be scaled or adjusted accordingly for each encoding experiment. We used Bitconverter's writing tool to generate pipetting commands for the epMotion automated liquid handler (Eppendorf) using a 96-well plate format for source and destination plates. The tool calculates the per-Guide transfer volume per population as the total donor cell volume (64 µl) divided by the number of edits per population, i.e. the number of Guides per population. It then computes the total volume demand per Guide culture across all populations and determines the number of source wells required to accommodate the demand within a user-defined per-well volume cap. A source volume summary table is produced, used for how much culture volume is required of each Guide strain. A fixed volume (16 µl) is used of the Writer for each population, and the total volume required is calculated as the number of populations x 16 µl.

Here, we describe exemplary programming of 96 CONCORD populations (one 96-well plate) on the epMotion. We grew up the Writer MG1655(T7RNAP) harbouring pLF177 to saturation overnight in 25 ml LB with chloramphenicol selecting for the base editor plasmid and IPTG inducing base editor expression. We also grew up 31 Guide cultures (Guide strains 1-7 & 9-32 used for SpCas9-CBE encoding) to saturation overnight in 10 ml LB with kanamycin selecting for the sgRNA plasmid and apramycin selecting for the F-plasmid. We collected the cells from each culture by centrifugation and washed the pellets once with 1 ml LB medium. We then resuspended the Writer in 2.5 ml MilliQ $H_2O$, and each Guide in 1 ml MilliQ $H_2O$. We distributed Writer and Guide cultures in a 96-well plate, each Guide culture into 4 wells with 210 µl per well, and the Writer culture into 10 wells with 210 µl per well, allowing for 50 µl dead volume. This 96-well plate functioned as source plate. We also provided an empty destination 96-well plate. We programmed pipetting of Guides into a combined 64 µl donor cell mixture in the destination plate and then the addition of 16µl of Writer culture to each mixture (4:1 ratio of donor:recipient). The donor cell mixture contains equal volumes of each Guide. For example, if population #1 requires editing at positions 5, 9, 12, 18, and 30, then 12.8 µl (64 µl total volume/5 Guides) of each of the corresponding Guide cultures is combined into one population mixture. Conjugation begins with the addition of the Writer cells. After addition of the Writer, 5 µl of each population mixture was immediately transferred into a new plate filled with 100 µl LB agar and once the spots were dried the plate was incubated at 37˚C for 1 h. After conjugative transfer, we resuspended cells in 200 µl LB medium with chloramphenicol and kanamycin selecting for recipient cells that received the sgRNA plasmid, and IPTG for base editor expression. After 6 h, 10 µl of the cultures were transferred into 190 µl of the same medium in a new plate. This was repeated twice more. After 24 h we incubated cultures for 5 min at 96˚C in a thermocycler, terminating the reactions, and stored them at -20˚C.

In total we encoded 45,689 bits of data in DNA, including 6x24 bits and 48x32 bits of sense-less data to probe the encoding limit, 75x31 bits and 84x27 bits in test datasets, 220x31 and 252x27 in Voyager datasets, and 832x31 in the Petroglyph dataset.

<u>Barcoding and pooling of written DNA data</u>

We designed 20 primers for PCR amplification and barcoding of the writing sequence with unique combinations of forward and reverse primers suitable for unique barcoding of 96 populations. The primers have the same binding site (forward or reverse) but distinct non-binding barcodes at the 5' ends. We chose 26-nucleotide barcode sequences from Oxford Nanopore Technologies' (ONT) Native Barcoding Kit 96 (SQK-NBD114.96, ONT), which are already optimised for subsequent demultiplexing. For PCR barcoding, 5 µl of OneTaq Quick-Load 2x Master Mix (New England Biolabs) was added to 1 µl of 1:10 diluted CONCORD encoding population mixture, 3 µl barcoded primer mix (forward and reverse primers, 1 µM) and topped up with nuclease free water to a total volume of 10 µl. The PCR reaction was run as per manufacturers protocol. Then, 5 µl of each PCR reaction was mixed into one combined pool, and purified with the DNA Clean & Concentrator-5 Kit (Zymo) following manufacturer's protocol. The pooled and purified barcoded amplicon pool was stored at -20°C.

<u>Reading DNA data by sequencing</u>

Amplicon pools were prepared for Nanopore sequencing using the Native Barcoding kit 24 (SQK-NBD114.24, ONT) following the manufacturer's protocol for sequencing of time-course and 6-bit codes directly programmed into the native *lacZ* locus. For all other encoded data we barcoded reactions by PCR as described above and prepared amplicons for Nanopore sequencing with the Ligation sequencing amplicons V14 kit (SQK-LSK114, ONT) following the manufacturer's protocol. 200 fmol of purified PCR amplicon pool was used as starting material for end-prep, directly followed by adapter ligation and finally eluted in 15 µl Elution Buffer. Sequencing was performed on MinION R10.4.1 flowcells (FLO-MIN114, ONT) on the MinION Mk1B sequencer (ONT) until ≥ 2 Gb of bases were called per pool of 96 populations, typically within less than 12 hours. Basecalling was conducted using the super-accurate basecalling model at default settings and basecalled reads were filtered at the default minimum overall read Q-Score of 10. Flow cells were washed using the flow cell wash kit (EXP-WSH004, ONT) as per manufacturer's protocol and re-used for subsequent sequencing.

<u>Sequencing read processing, custom denoising and calculation of editing rates</u>

Sequencing reads were filtered by length (selecting for reads between 1500-2500 bases), demultiplexed by their terminal barcodes and aligned to a reference unedited FASTA sequence (all target bases in 0 state) on the EPI2ME Desktop (V5.2.1) using the wf-alignment workflow (V1.2.5) with default settings. After alignment, the .bam and .bai files were analyzed using our custom denoising script (**Supplementary Data 21**): The 'Sam2Tsv' function of the *JVARKIT* program was used to print the SAM alignments, with each line corresponding to an individual base of a read in the alignment. *Perl* was used to convert the Q-Scores from ASCII to numerical integers. For each read, we analysed target positions as follows: reads were included if the target position was aligned, as indicated by an M operation in the CIGAR string, and if the position-specific read quality is above Q-Score 20. The total number of sequenced "A/T/G/C" bases above the Q-Score threshold was listed for each target locus, and the editing rate was calculated as ratio T/(A+T+G+C) for (C→T) substitutions and G/(A+T+G+C) for (A→G) substitutions. We filtered out approximately 10% of reads during the denoising step. To assess the effect of sequencing depth on data recovery we created random subsamples of 10, 100, or 1000 reads per population with a custom python script **(Supplementary Data 21)** and calculated editing rates as described. Consequently, a minimum sequencing depth of 1000 reads per population after denoising was set as the requirement for base editing calculations and sequencing was repeated for populations with below threshold depth. For each population, we determined the editing rate for one canonical target base at all target positions (one editing rate per encoded bit). The canonical base was identified in populations with single Guides as the base in the protospacer sequence with the highest editing rate. The final output is a matrix of editing rates with one editing rate per position for each population.

Decoding editing rates to binary data and conversion to digital data

First, we transformed editing rates (x) by shifted double log transformation: $y=\ln(\ln(1+x)+1)$. We decoded transformed editing rates to binary data by applying thresholding and clustering methods.

For thresholding, we first used a test dataset to train thresholds for both SpCas9-CBE and SpCas9-ABE in order to distinguish programmed "0"s from programmed "1"s. To this end we modelled the class-conditional distributions of the transformed editing rates for negatives (y=0) and positives (y=1) as Gaussian distributions with parameters μ0, σ0, μ1, and σ1, estimated as the empirical mean and standard deviation of each class. The decision threshold was set as the intersection of the two Gaussian probability density functions p(x), computed as the solution to $p_0(x) = p_1(x)$. This was computed globally using editing rates across all positions (global threshold) or separately for each position (local thresholds). Class priors are recorded but not used in the intersection criterion. Computed thresholds are listed in **Supplementary Data 4**. For threshold-based decoding we then decoded editing rates > threshold as "1" and editing rates ≤ threshold as "0". This was either done for all editing rates independent of encoding position using the global threshold or per position using local thresholds. Accuracy of the decoding was calculated as the fraction of correctly decoded bits, computed for all bits (global threshold decoding) or per position (local threshold decoding).

For decoding by clustering, we performed six unsupervised machine-learning algorithms (GMM, DBSCAN, K-Means, HDBSCAN, OPTICS, Mean-Shift) with two target groups and mapped clusters to binary labels by assigning the cluster with the higher transformed mean to "edited" (1) and the lower to "unedited" (0). Points labelled as noise were assigned by proximity to the two-cluster means, and if ≤1 non-noise cluster was found we used a median split. Analogous to threshold decoding we performed clustering on all editing rates (global clustering) and per position (local clustering). Decoding accuracies were calculated as above.

In the final CONCORD workflow, we combined local clustering decoding with GMM, HDBSCAN, and K-Means by majority vote on a per-bit basis. For each provided 0/1 in the binary matrix (populations x positions), we count votes for 1 and 0 across files and assign the majority. We then concatenated the binary matrix into one binary string and converted the string back to digital data using the same character encoding standard used for encoding with the Bitconverter data conversion tool.

For redundantly encoded data with SpCas9-CBE and SpCas9-ABE we added an additional decision certainty layer for data recovery. For discrepancies in converted digital data, we first assessed individual binary predictions from local clustering (GMM, HDBSCAN, K-Means) for both Writers and computed the median value. For uncertain clustering results (median = 0.5), we used a threshold-based score to decide on the bit-value with more certain experimental base editing data. Essentially, we assess the distance of the editing rate to the respective previously defined local and global Gaussian thresholds and prioritise the experiment with greater distance. To this end we computed posteriors P(edited|x) and local likelihood ratios $LLR = \log f_1(x) - \log f_0(x)$ for editing rates of uncertain bits for both Writers. For each Writer, the two LLRs were combined as a weighted sum and added to the log prior odds, yielding a combined logit and posterior via the logistic function. Final calls used a probability threshold with a symmetric gray zone: $p < (\tau - \gamma) \rightarrow$ "unedited", $p > (\tau + \gamma) \rightarrow$ "edited", otherwise "ambiguous".

Custom python scripts used for threshold computing, clustering, majority clustering voting, and threshold-based scoring are available **(Supplementary Data 21).**

Random access

We demonstrated random access of data by recovering six selected words from the Voyager text encoded by SpCas9-CBE and SpCas9-ABE and stored as three amplicon pools each. We first identified the populations in which the relevant bits were encoded. We then identified the barcodes attached to amplicons from the populations of interest. We ordered primers matching the barcodes and amplified writing sequences of interest from the pools by PCR with Q5® Hot Start High-Fidelity 2X Master Mix (New England Biolabs) following the manufacturer's protocol with 1 ng of amplicon pool as template input and 35 cycles. We analysed 5 µl of the PCR reactions on a 1% agarose gel, and processed the rest for Nanopore sequencing as described above. Following sequencing, we determined editing rates at all positions and decoded the editing rates to binary data by majority vote local clustering (GMM, HDBSCAN, K-Means) as described above.

Materials and Data availability

All materials and data are available upon request. Sequences of all relevant plasmids and synthetic writing sequences are provided as Genbank files in **Supplementary Data 8-19**. Editing rates data are available in **Supplementary Data 3 and 6**. Decoding results are available in **Supplementary Data 5 and 7**. Code is available in **Supplementary Data 21**.

### Extended Data Figures

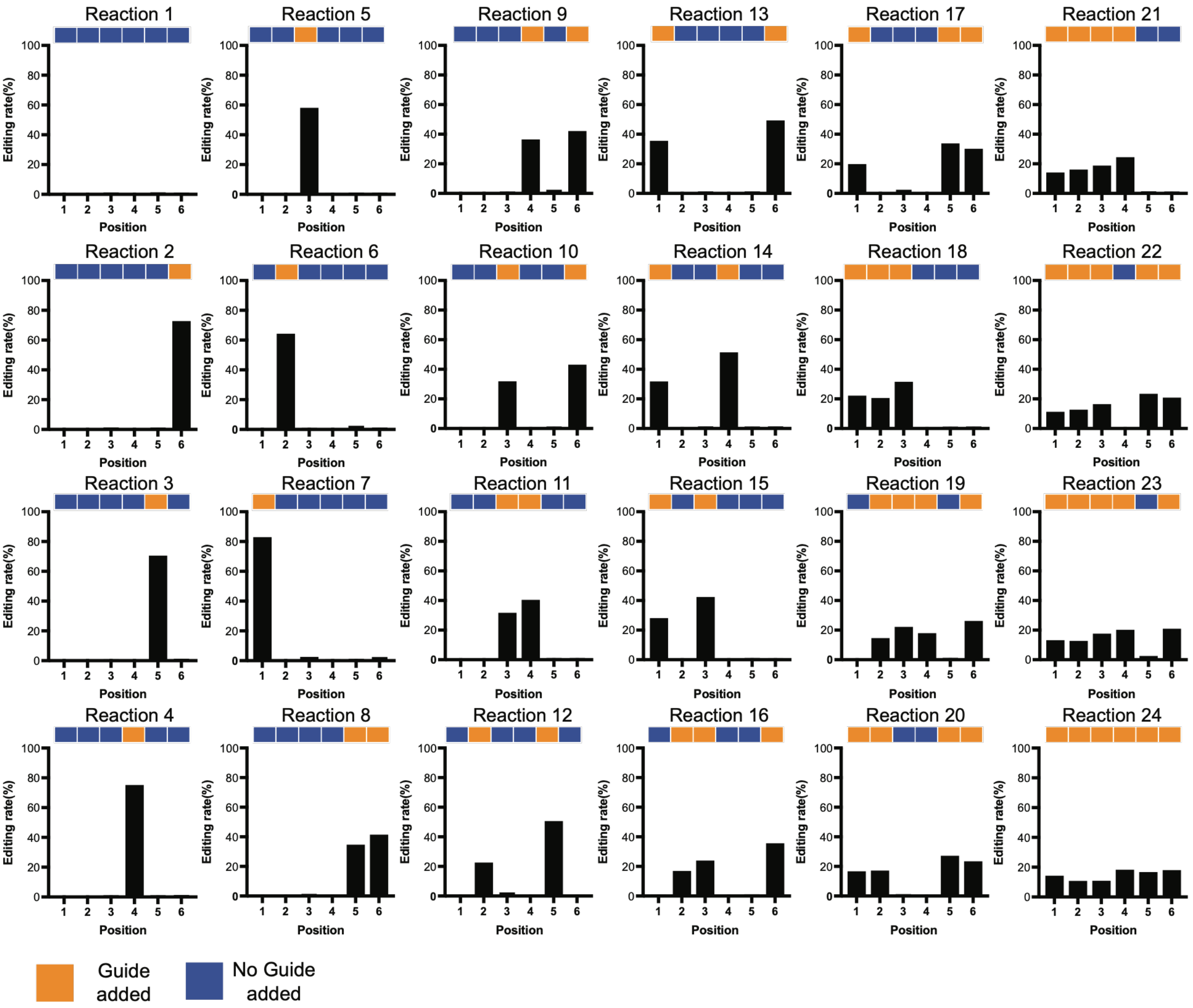


**Extended Data Figure 1: Programmed patterns and editing rates for 24 populations with up to six Guides added per population.** Code programming schemes are depicted above each graph. Up to six genomic positions in the lacZ gene were targeted. Reactions were terminated 24 hours after conjugation and editing rates were recorded at six designated target cytosines for each population. Each encoding was performed in a single replicate and editing rates recorded after 24h are shown.

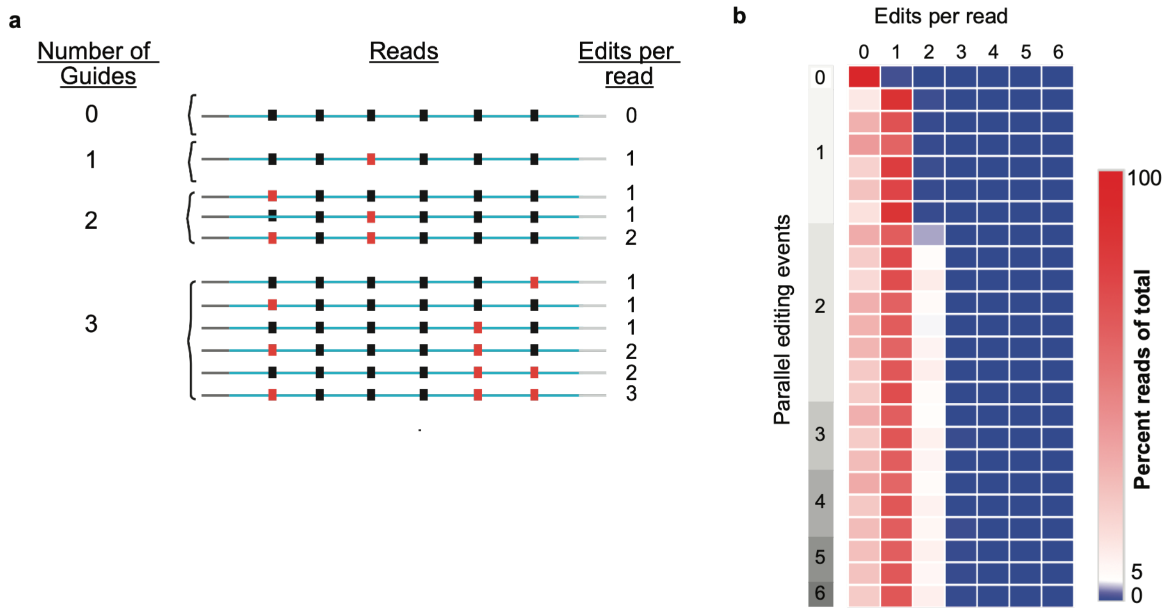


**Extended Data Figure 2: Assessment of multiplexed base edits within one Writer cell. (A)** Schematic of theoretical editing events in recipient cells in a population with a Writer strain and multiple distinct Guide strains. All Guides carry a non-transferrable F-plasmid, which makes the recipient Writer cells susceptible for continuous horizontal gene transfers, as it remains F-. **(B)** Percentage of reads with multiple target positions edited. To assess editing at multiple positions in the same cell we performed a special read filtering after sequencing, selecting only reads with high quality reads at all six target positions. Percentages of reads with 0-6 edits within those reads were recorded for all 24 populations.

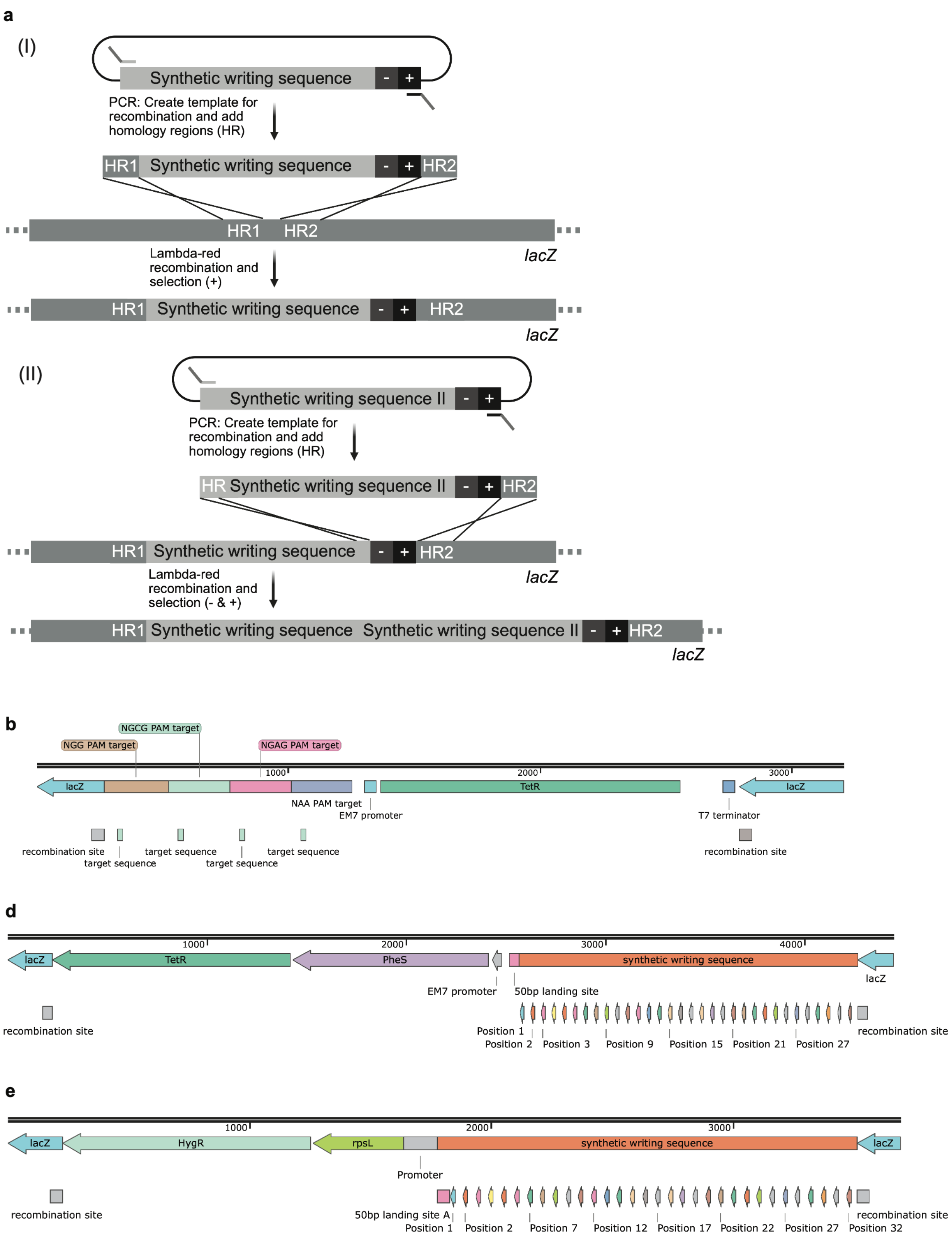


**Extended Data Figure 3: Schematics of synthetic writing sequences for CONCORD experiments and integration into the *E. coli* genome. (a)** (I) A designed writing sequence was synthesized and cloned into an intermediate plasmid and then amplified coupled to a double selection cassette consisting of a negative (-) and positive (+) selection maker to create a template for homologous recombination. Short, 50bp homology regions were introduced in the amplification at the ends, homologous to a target in the lacZ gene (HR1, HR2). Amplicons were electroporated into recipient MG1655 cells expressing components for lambda-red recombination. The positive selection marker was used to select for cells with successful genomic integration of the construct. (II) The negative selection marker facilitates simple extension of the writing sequence in adjacent integrations. This integration strategy has been described previously[2]. **(b)** Map of gLF3456 integrated genomically at the MG1655(T7RNAP) *lacZ* locus. A ˜1kb cassette with four repeats of one protospacer that can be targeted with the same sgRNA. Each repeat is flanked by a distinct PAM sequence, creating four target positions. The cassette was coupled to *tetR* (+) for positive selection after integration. **(c)** Map of gLF013 integrated genomically at the MG1655(T7RNAP) *lacZ* locus. The ˜1.7kb cassette with 32 protospacers followed by NGG PAMs and 30 bp spacing sequences in between protospacers, was coupled to a mutated *pheS** (-) and *tetR* (+). **(d)** Map of gLF014 integrated genomically at the MG1655(T7RNAP) *lacZ* locus. The ˜1.7kb cassette with 32 protospacers followed by NAA PAMs and 30 bp spacing sequences in between protospacers, was coupled to *rpsL* (-) and *hygR* (+) for selection. All sequences are also included as Genbank files in **Supplementary Data 17-19**.

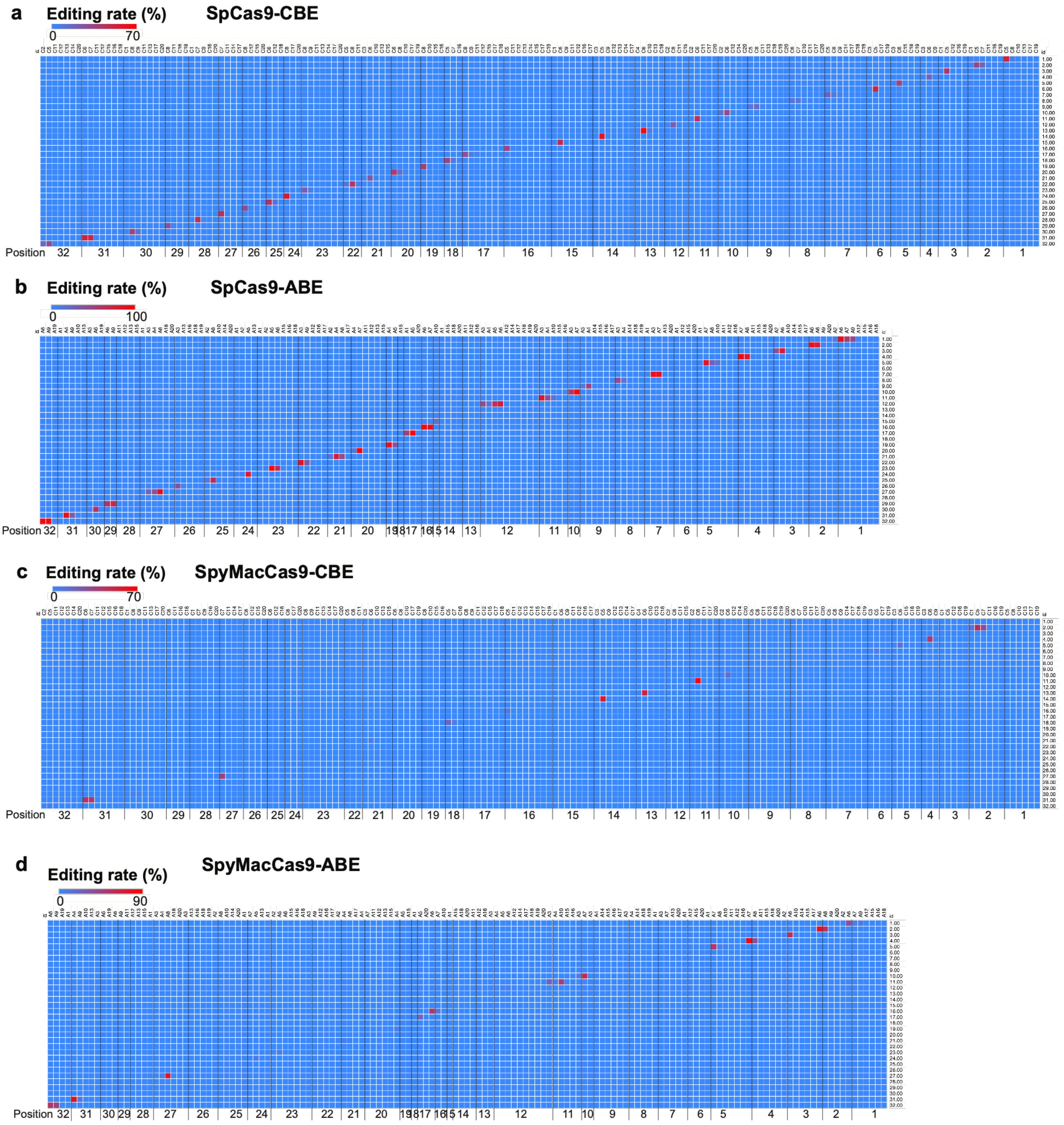


**Extended Data Figure 4: Target specificity of 32 sgRNAs assessed at all 32 protospacers present in synthetic writing cassettes gLF013 and gLF014.** Base editing rates at all cytosines and adenines were assessed in populations of Guide strains 1-32 with SpCas9-CBE **(a)**, SpCas9-ABE **(b)**, SpyMacCas9-CBE **(c)**, SpyMacCas9-ABE **(d)** after read filtering. Editing rate scale bars are adjusted to the maximum editing rates recorded for each Writer to resolve low or off-target editing as well as possible. Each experiment was performed in biological duplicates, and means are shown in the heatmaps.

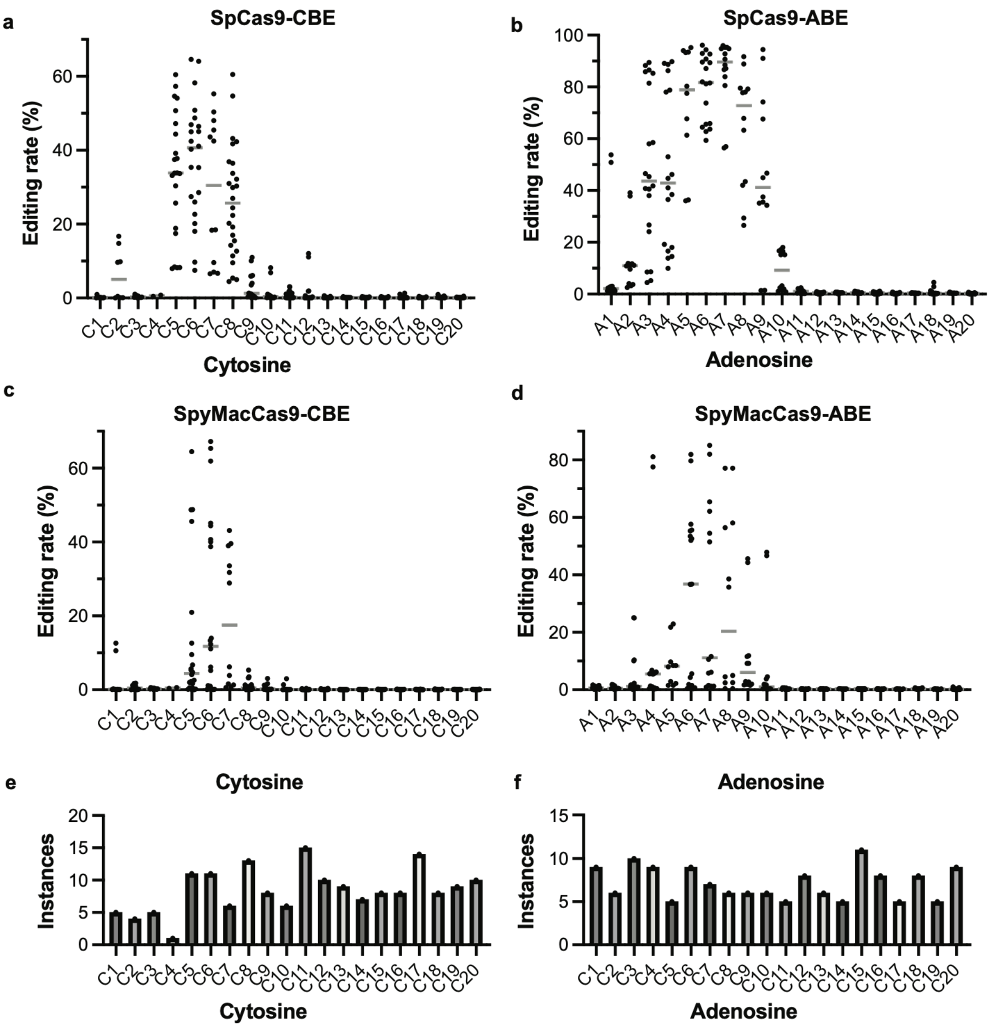


**Extended Data Figure 5: Editing patterns of four orthogonal base editors.** Base editing rates at all cytosines or adenosines present in a protospacer were recorded across the 32 target positions in the synthetic writing sequence from biological duplicate experiments of each Writer with each Guide. All individual recorded editing rates are shown, grouped by position of the base within the 20-bp protospacer counting from the PAM-opposite end (PAM = 21-23). Medians are shown as grey bars. Cytosine editing rates were recorded for SpCas9-CBE **(a)** and SpyMacCas9-CBE **(c)**. Adenine editing rates were recorded for SpCas9-ABE **(b)** and SpyMacCas9-ABE **(d)**. Number of cytosines **(e)** and adenines **(f)** present assessed for each relative protospacer position across the 32 protospacers.

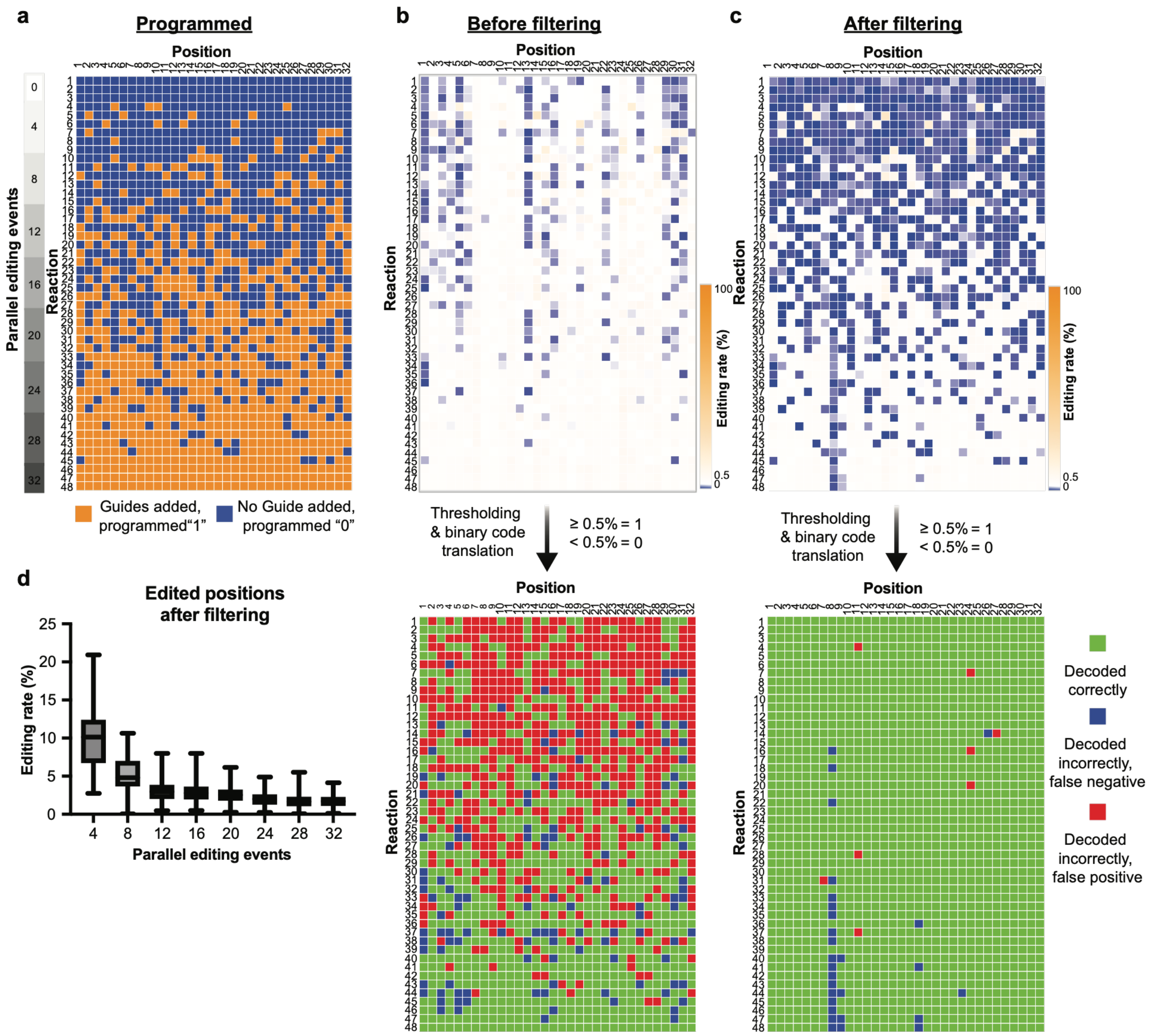


**Extended Data Figure 6: Position-specific read filtering is crucial for distinguishing programmed "1"s from "0"s when scaling up the number of parallel editing events. (a)** Programmed pattern of 48 populations to probe the detection limit of parallel base editing events with SpCas9-CBE. Each CONCORD experiment was performed in a single biological replicate **(b)** Editing rates at target positions before position-specific read filtering and decoding result using a 0.5% threshold. Green for sites correctly decoded as programmed, blue for sites falsely decoded as "0" while programmed "1", red for sites falsely decoded as "1" while programmed "0". **(c)** Editing rates recorded at target positions after position-specific read filtering and decoding result using a 0.5% threshold. **(d)** Editing rates at programmed "1"s across the 48 populations grouped by number of parallel editing events before and after position-specific read filtering. Standard box plots with median marked.

**a**

| Binary | Index (Decimal) | Character | Binary | Index (Decimal) | Character |
|---|---|---|---|---|---|
| 000000 | 0 | (space) | 100000 | 32 | 5 |
| 000001 | 1 | A | 100001 | 33 | 6 |
| 000010 | 2 | B | 100010 | 34 | 7 |
| 000011 | 3 | C | 100011 | 35 | 8 |
| 000100 | 4 | D | 100100 | 36 | 9 |
| 000101 | 5 | E | 100101 | 37 | . |
| 000110 | 6 | F | 100110 | 38 | , |
| 000111 | 7 | G | 100111 | 39 | ( |
| 001000 | 8 | H | 101000 | 40 | ) |
| 001001 | 9 | I | 101001 | 41 | + |
| 001010 | 10 | J | 101010 | 42 | - |
| 001011 | 11 | K | 101011 | 43 | * |
| 001100 | 12 | L | 101100 | 44 | / |
| 001101 | 13 | M | 101101 | 45 | = |
| 001110 | 14 | N | 101110 | 46 | ! |
| 001111 | 15 | O | 101111 | 47 | : |
| 010000 | 16 | P | 110000 | 48 | % |
| 010001 | 17 | Q | 110001 | 49 | " |
| 010010 | 18 | R | 110010 | 50 | # |
| 010011 | 19 | S | 110011 | 51 | @ |
| 010100 | 20 | T | 110100 | 52 | ? |
| 010101 | 21 | U | 110101 | 53 | & |
| 010110 | 22 | V | 110110 | 54 | (Image) |
| 010111 | 23 | W | 110111 | 55 | Unused |
| 011000 | 24 | X | 111000 | 56 | Unused |
| 011001 | 25 | Y | 111001 | 57 | Unused |
| 011010 | 26 | Z | 111010 | 58 | Unused |
| 011011 | 27 | 0 | 111011 | 59 | Unused |
| 011100 | 28 | 1 | 111100 | 60 | Unused |
| 011101 | 29 | 2 | 111101 | 61 | Unused |
| 011110 | 30 | 3 | 111110 | 62 | Unused |
| 011111 | 31 | 4 | 111111 | 63 | Unused |

**b**

| Binary | Index (Decimal) | 8-bit Equivalent (0–255) | Approx. brightnes |
|---|---|---|---|
| 0000 | 0 | 0 | 0.00% |
| 0001 | 1 | 17 | 6.70% |
| 0010 | 2 | 34 | 13.30% |
| 0011 | 3 | 51 | 20.00% |
| 0100 | 4 | 68 | 26.70% |
| 0101 | 5 | 85 | 33.30% |
| 0110 | 6 | 102 | 40.00% |
| 0111 | 7 | 119 | 46.70% |
| 1000 | 8 | 136 | 53.30% |
| 1001 | 9 | 153 | 60.00% |
| 1010 | 10 | 170 | 66.70% |
| 1011 | 11 | 187 | 73.30% |
| 1100 | 12 | 204 | 80.00% |
| 1101 | 13 | 221 | 86.70% |
| 1110 | 14 | 238 | 93.30% |
| 1111 | 15 | 255 | 100.00% |

**c**

| Binary | Index (Decimal) | R (0/255) | G (0/255) | B (0/255) | Color Hex | Common Name |
|---|---|---|---|---|---|---|
| 000 | 0 | 0 | 0 | 0 | #000000 | Black |
| 001 | 1 | 0 | 0 | 255 | #0000FF | Blue |
| 010 | 2 | 0 | 255 | 0 | #00FF00 | Green |
| 011 | 3 | 0 | 255 | 255 | #00FFFF | Cyan |
| 100 | 4 | 255 | 0 | 0 | #FF0000 | Red |
| 101 | 5 | 255 | 0 | 255 | #FF00FF | Magenta |
| 110 | 6 | 255 | 255 | 0 | #FFFF00 | Yellow |
| 111 | 7 | 255 | 255 | 255 | #FFFFFF | White |

**Extended Data Figure 7: Character encoding standards used in this study. (a)** 6-bit LNS code used to encode part of the test datasets, the Voyager text, as well as introductory text in the Petroglyph dataset. **(b)** 4-bit grey-scale code used to encode the Petroglyph image. **(c)** 3-bit RGB code used to encode the rainbow image in the test dataset.

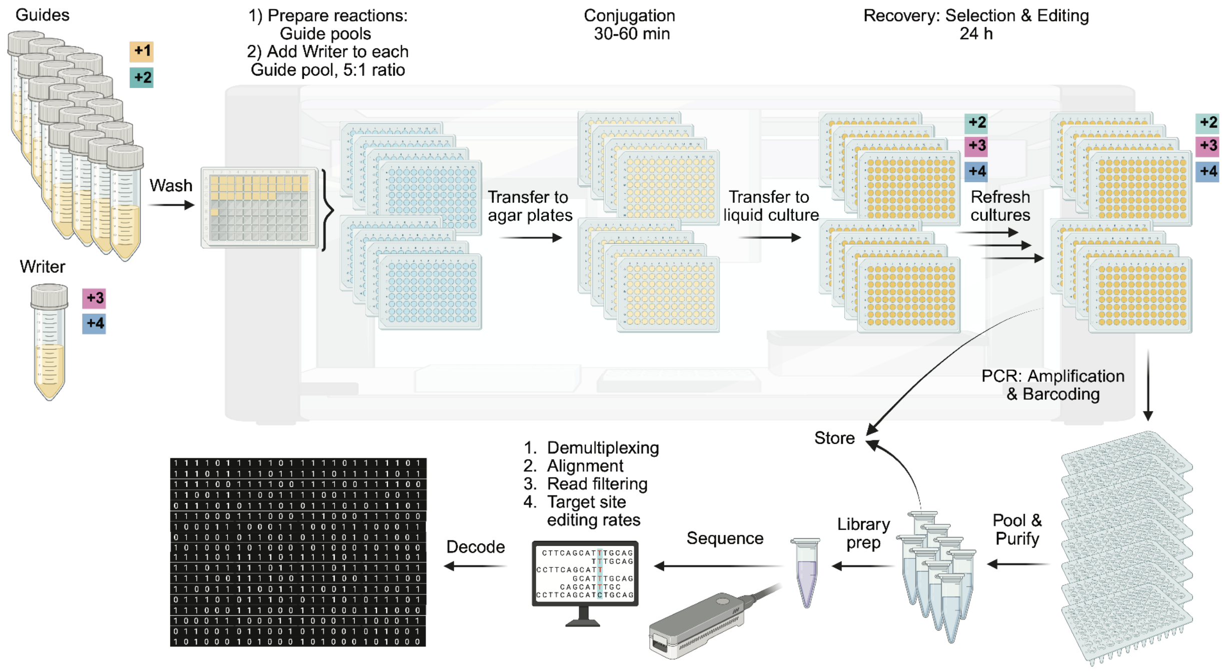


**Extended Data Figure 8: Schematic of CONCORD workflow.** Guide and Writer cell cultures are grown up over night: Guide strains with chloramphenicol (+1) and apramycin (+2) selection for the sgRNA plasmid and F-plasmid, the Writer culture with kanamycin (+3) selection for the base editor plasmid and IPTG (+4) for induction of base editor expression. Following one round of washing to remove antibiotics the cultures are distributed in a multi-well plate. Reactions are programmed on a liquid handler, combining selected Guides into pools for each population, and adding the Writer to initiale conjugation. For conjugation, 5 µl of each population is transferred into 96-well plates filled with agar, and incubated at 37˚C for 1 hour to facilitate horizontal gene transfer. Populations are then resuspended in 200 µl liquid medium with antibiotics (+2, +3) selecting for Writers that received the sgRNA plasmid and IPTG (+4) to maintain base editor expression. Cultures are refreshed 2-3 times during the recovery phase while shaking at 37˚C to increase base editing rates. Reactions are terminated after 24 hours and can be stored as cells or as amplicon pools. To recover encoded information amplicons are sequenced followed by data processing and decoding. The encoding process is highly parallelizable and scalable and requires minimal volumes of inexpensive cell culture inputs.

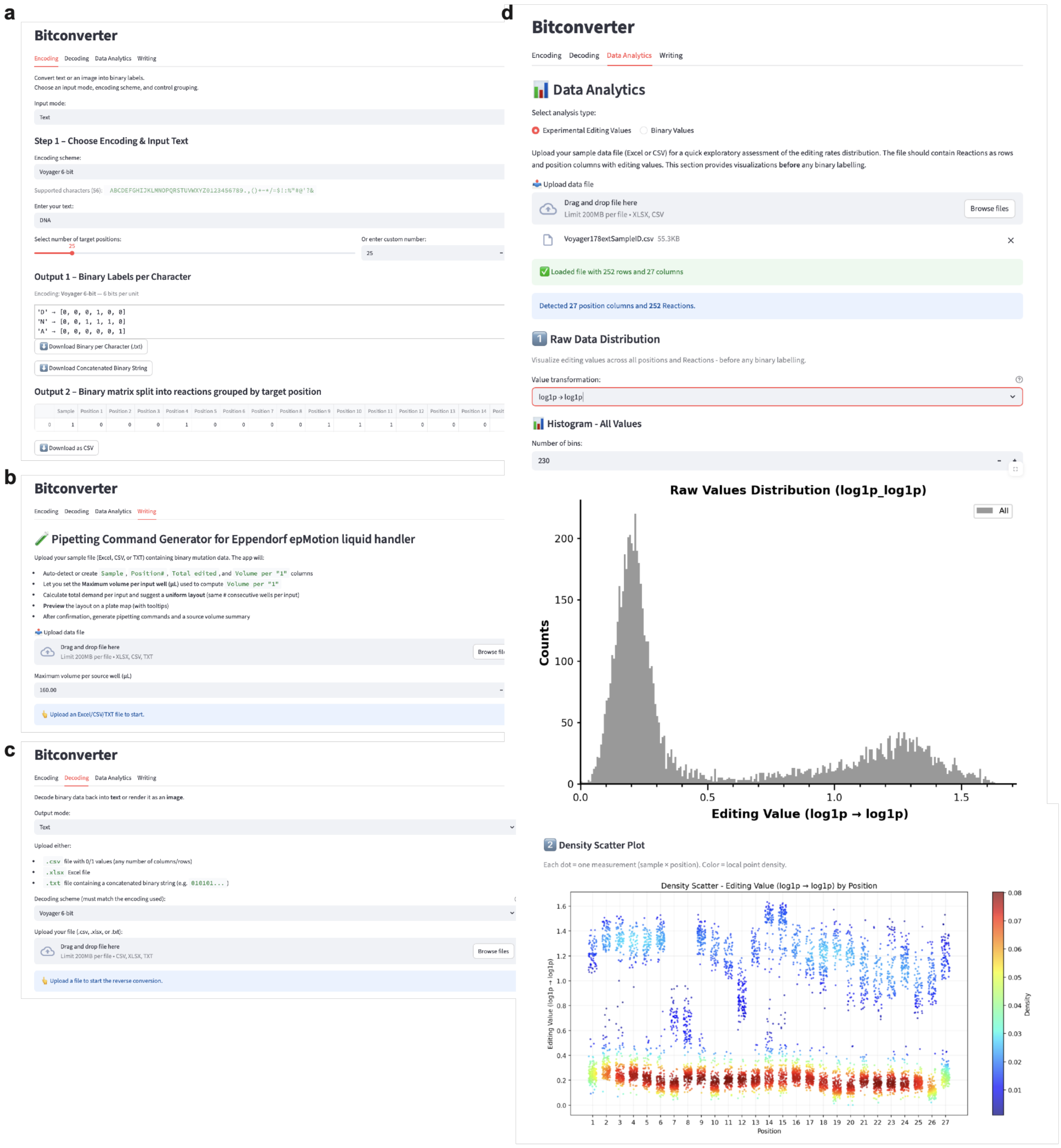


**Extended Data Figure 9: Bitconverter is a freely available, browser-based web application with tools to support CONCORD data encoding and decoding.** Bitconverter bridges the gap between digital information encoding and the practical wet-lab requirements for DNA data encoding. Bitconverter is implemented in Python using the Streamlit framework and is deployed on Hugging Face Spaces (https://huggingface.co/spaces/EngBioNUS/bitconverter). **(a)** Encoding tool. Bitconverter supports four character-to-binary encoding schemes, each producing a fixed number of bits per encoding unit. It then converts the binary code into programmed populations adjusting to the number of target positions utilized by the Writer of choice. Bitconverter also converts raster images into binary matrices using two quantization modes. In 1-bit (black and white) mode, images are converted to grayscale, resized to a user-specified width while preserving aspect ratio, and thresholded at a user-defined intensity cutoff (default 128). In 4-bit grayscale mode, 8-bit grayscale pixel values (0-255) are uniformly quantized to 16 levels (0-15). The complete list of populations can be downloaded as binary matrix grouped by target positions. **(b)** Writing tool. Bitconverter includes a pipetting command generator for the Eppendorf epMotion automated liquid handler using a 96-well plate format for source and destination plates. The tool calculates the per-population transfer volumes for Guides, computes the total volume demand per source input across all samples and determines the number of source wells required to accommodate the demand within a user-defined per-well volume cap. A source volume summary table is produced as well as a source plate layout. **(c)** To assist users in selecting appropriate binarization strategies from continuous editing rate data, Bitconverter provides an exploratory data analytics module. Users upload tabular data (CSV or Excel) containing editing rates across samples and positions. The module automatically detects numeric position columns and offers three value transformations: raw linear values, shifted log-transformation (ln(1+x)), and shifted double log transformation (ln(ln(1 + x)+1)). Two interactive visualizations are generated: a histogram of all unlabelled values, and a density scatter plot of editing values by target position coloured by kernel density estimation. **(d)** Conversion tool. Binary code generated after decoding of editing rates can be converted back to characters or image with the conversion tool. Bitconverter supports four binary-to-character decoding schemes for text, 1-bit (black-and-white) and 4-bit grey scale modes for image decoding. A decoded image is rendered with nearest-neighbour scaling for crisp pixel visualization and exported as PNG.

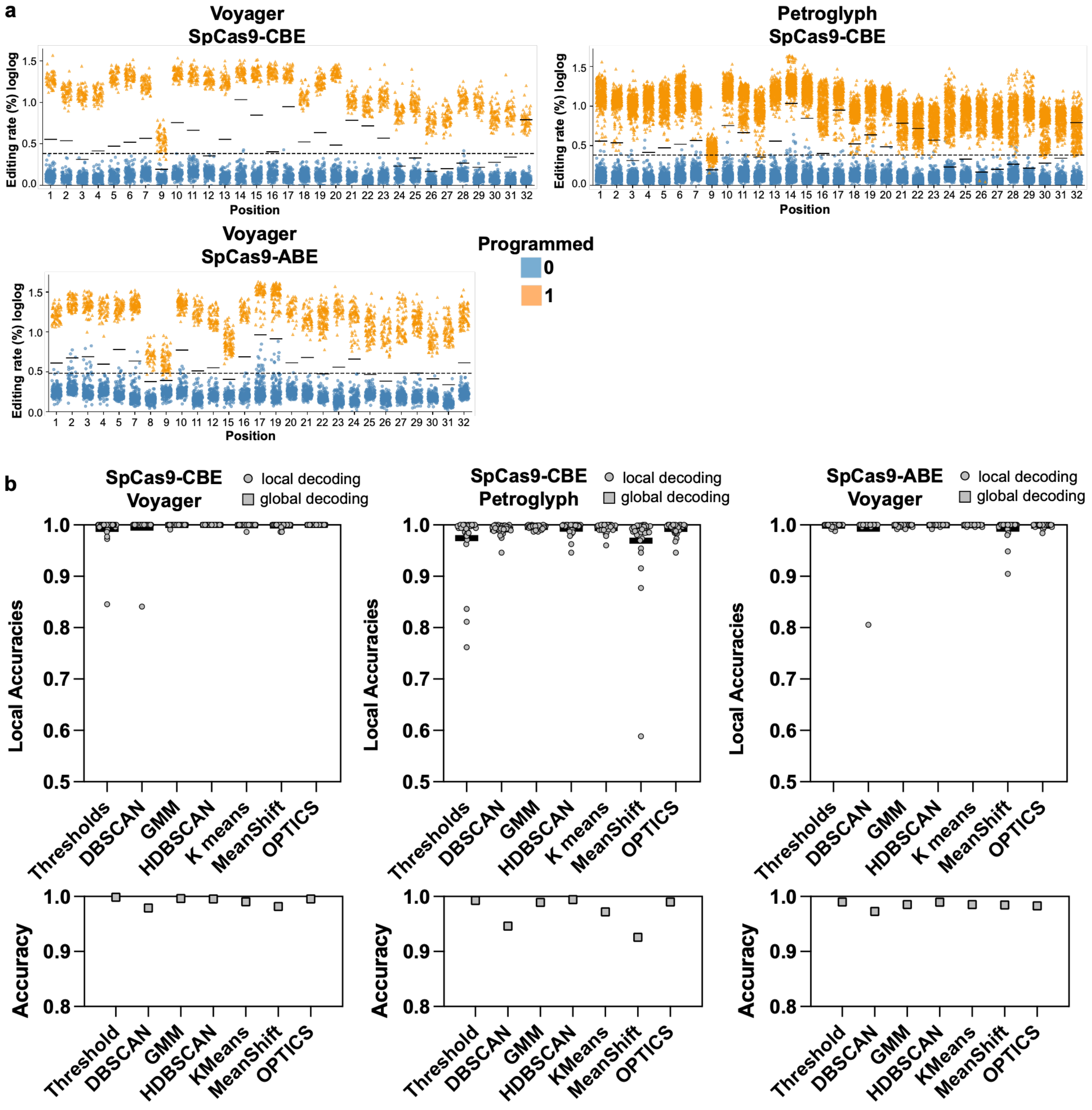


**Extended Data Figure 10: Decoding Voyager and Petroglyph datasets ‘blindly’. (a)** Double log transformed editing rates grouped by target position for three independent CONCORD encodings. Data points are labelled by their intended programmed state (programmed “1”s orange, programmed “0”s blue). Thresholds learned on the test dataset for SpCas9-CBE and SpCas9-ABE are shown (dashed line for global threshold, solid black lines for local thresholds. Positions refer to the 32 sites on the NGG cassette. **(b)** Accuracies of decoding each dataset by supervised learning (thresholds) as well as unsupervised learning (clustering) strategies. Clustering algorithms include DBSCAN, HDBSCAN, GMM, KMeans, MeanShift, and OPTICS. Accuracy refers to the fraction of correctly decoded bits. Data sets were decoded by position (local) or as one data set (global).

**a**

THIS VOYAGER SPACECRAFT WAS CONSTRUCTED BY THE UNITED STATES OF AMERICA. WE ARE A[81]
COMMUNITY OF 240 MILLION HUMAN BEINGS AMONG THE MORE THAN 4 BILLION WHO INHABIT THE[165]
PLANET EARTH. WE HUMAN BEINGS ARE STILL DIVIDED INTO NATION STATES, BUT THESE STATES[250]
ARE RAPIDLY BECOMING A GLOBAL CIVILIZATION. WE CAST THIS MESSAGE INTO THE COSMOS ... IT[338]
IS LIKELY TO SURVIVE A BILLION YEARS INTO OUR FUTURE, WHEN OUR CIVILIZATION IS[417]
PROFOUNDLY ALTERED AND THE SURFACE OF THE EARTH MAY BE VASTLY CHANGED. OF THE 200[499]
BILLION STARS IN THE MILKY WAY GALAXY, SOME PERHAPS MANY MAY HAVE INHABITED[579]
PLANETS AND SPACE FARING CIVILIZATIONS. IF ONE SUCH CIVILIZATION INTERCEPTS VOYAGER[663]
AND CAN UNDERSTAND THESE RECORDED CONTENTS, HERE IS OUR MESSAGE: THIS IS A[738]
PRESENT FROM A SMALL DISTANT WORLD, A TOKEN OF OUR SOUNDS, OUR SCIENCE, OUR[814]
IMAGES, OUR MUSIC, OUR THOUGHTS, AND OUR FEELINGS. WE ARE ATTEMPTING TO SURVIVE[894]
OUR TIME SO WE MAY LIVE INTO YOURS. WE HOPE SOME DAY, HAVING SOLVED THE PROBLEMS[975]
WE FACE, TO JOIN A COMMUNITY OF GALACTIC CIVILIZATIONS. THIS RECORD REPRESENTS OUR[1058]
HOPE AND OUR DETERMINATION AND OUR GOODWILL IN A VAST AND AWESOME UNIVERSE.[1134]

**Acc** 1.000
**FP** 0
**FN** 0

**b**

THIS VOYAGER SPACECRAFT WAS CONSTRUCTED BY THE UNITED STATES OF AMERICA. WE ARE A [81]
COMMUNITY OF 240 MILLION HUMAN BEINGS AMONG THE MORE THAN 4 BILLION WHO INHABIT THE [165]
PLANET EARTH. WE HUMAN BEINGS ARE STILL DIVIDED INTO NATION STA**U**ES, BUT THESE STATES[250]
ARE RAPIDLY BECOMING A GLOBAL CIVILIZATION. WE CAST THIS MESSAGE**H**INTO THE COSMOS ... IT[338]
IS LIKELY TO SURVIVE A BILLION YEARS INTO OUR FUTURE, WHEN OUR CIVILIZATION IS [417]
PROFOUNDLY ALTERED AND THE SURFACE OF THE EARTH MAY BE VASTLY CHANGED. OF THE 200 [499]
BILLION STARS IN THE MILKY WAY GALAXY, SOME PERHAPS MANY MAY HAVE INHABITED [579]
PLANETS AND SPACE FARING CIVILIZATIONS. IF ONE SUCH CIVILIZATION INTERCEPTS VOYAGER [663]
AND CAN UNDERSTAND THESE RECORDED CONTENTS, HERE IS OUR MESSAGE: THIS IS A [738]
PRESENT FROM A SMALL DISTANT WORLD, A TOKEN OF OUR SOUNDS, OUR SCIENCE, OUR [814]
IMAGES, OUR MUSIC, OUR THOUGHTS, AND OUR FEELINGS. WE ARE ATTEMPTING TO SURVIVE [894]
OUR TIME SO WE MAY LIVE INTO YOURS. WE HOPE SOME DAY, HAVING SOLVED THE PROBLEMS [975]
WE**A**FACE, TO JOIN A COMMUNITY OF GALACTIC CIVILIZATIONS. THIS RECORD REPRESENTS OUR[1058]
HOPE AND OUR DETERMINATION AND OUR GOODWILL IN A VAST AND AWESOME UNIVE**P**SE. [1134]

**Acc** 0.999
**FP** 3
**FN** 1

**c**

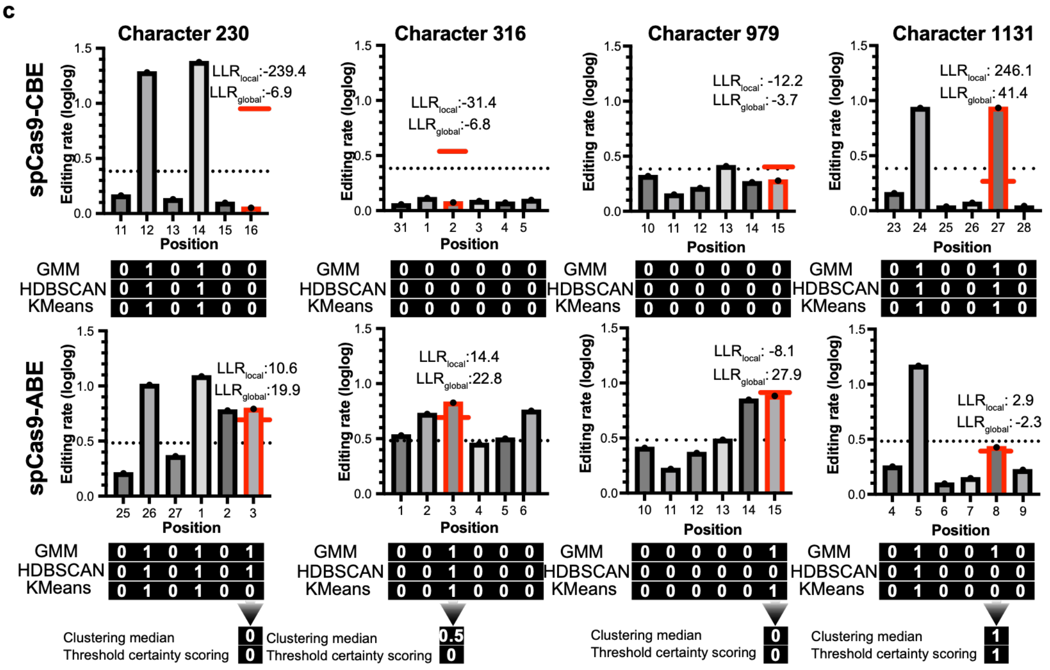


**Extended Data Figure 11: Voyager text recovered and discrepancies in the recovered text.** **(a)** Voyager text recovered after DNA encoding with SpCas9-CBE, and decoding of editing rates by majority vote local clustering with GMM, HDBSCAN, and KMeans algorithms. The

Voyager greeting contains a total of 1134 characters including spaces. Character numbers are indicated in superscript at the end of each line. Incorrectly decoded characters are highlighted in red. Accuracy (Acc) of decoding as well as number of false positives (FP) and false negatives (FN) are listed. **(b)** Voyager text recovered after DNA encoding with SpCas9-ABE, and decoding of editing rates by majority vote local clustering with GMM, HDBSCAN, and KMeans algorithms. **(c)** Strategy for solving discrepancies in redundancy encoded data. The median over 6 local clustering predictions from GMM, HDBSCAN, and KMeans was formed and certainty scores (LLR) were computed based on distance of editing rates from global or local thresholds at the respective positions. In case of uncertain clustering (median of 0.5), a threshold-based certainty scoring approach was followed taking all four certainty scores into account for an uncertain bit. Plots show single value editing rates since all encodings were performed in single biological replicates.

**a**

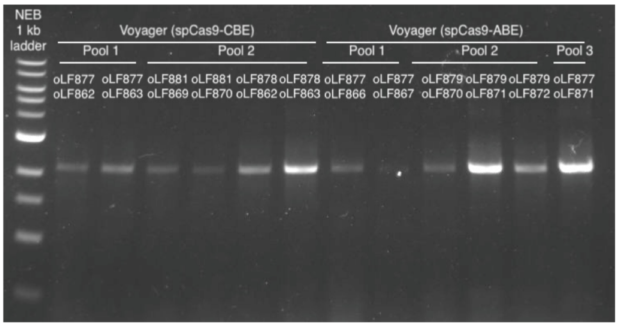


**b** THIS VOYAGER SPACECRAFT WAS CONSTRUCTED BY THE UNITED STATES OF AMERICA. WE ARE A COMMUNITY OF 240 MILLION HUMAN BEINGS AMONG THE MORE THAN 4 BILLION WHO INHABIT THE PLANET EARTH. WE HUMAN BEINGS ARE STILL DIVIDED INTO NATION STATES, BUT THESE STATES ARE RAPIDLY BECOMING A GLOBAL CIVILIZATION. WE CAST THIS MESSAGE INTO THE COSMOS ... IT IS LIKELY TO SURVIVE A BILLION YEARS INTO OUR FUTURE, WHEN OUR CIVILIZATION IS PROFOUNDLY ALTERED AND THE SURFACE OF THE EARTH MAY BE VASTLY CHANGED. OF THE 200 BILLION STARS IN THE MILKY WAY GALAXY, SOME PERHAPS MANY MAY HAVE INHABITED PLANETS AND SPACE FARING CIVILIZATIONS. IF ONE SUCH CIVILIZATION INTERCEPTS VOYAGER AND CAN UNDERSTAND THESE RECORDED CONTENTS, HERE IS OUR MESSAGE: THIS IS A PRESENT FROM A SMALL DISTANT WORLD, A TOKEN OF OUR SOUNDS, OUR SCIENCE, OUR IMAGES, OUR MUSIC, OUR THOUGHTS, AND OUR FEELINGS. WE ARE ATTEMPTING TO SURVIVE OUR TIME SO WE MAY LIVE INTO YOURS. WE HOPE SOME DAY, HAVING SOLVED THE PROBLEMS WE FACE, TO JOIN A COMMUNITY OF GALACTIC CIVILIZATIONS. THIS RECORD REPRESENTS OUR HOPE AND OUR DETERMINATION AND OUR GOODWILL IN A VAST AND AWESOME UNIVERSE.

**c**

| Sample | Barcode oligos | Binary code | Decoding result | Decoding metrics |
|---|---|---|---|---|
| Pool 1 | oLF877/862<br>oLF877/863 | 00000100111000000000000100001010<br>01001001110000111010011000000000 | AN BEINGS | **Acc:** 1.000; **FP:** 0; **FN:** 0 |
| Pool 1 | oLF877/866<br>oLF877/867 | 010111000101000000001000010<br>101001101000001001110000000 | WE HUMAN | **Acc:** 1.000; **FP:** 0; **FN:** 0 |
| Pool 2 | oLF881/869<br>oLF881/870 | 00010000101000000001011000000101<br>00110101000011000110010000000000 | E VASTLY | **Acc:** 1.000; **FP:** 0; **FN:** 0 |
| Pool 2 | oLF878/862<br>oLF878/863 | 00111001010000000000001100100100<br>01111001101000000000000100000001 | NT FROM A | **Acc:** 1.000; **FP:** 0; **FN:** 0 |
| Pool 2 | oLF879/870<br>oLF879/871<br>oLF879/872 | 000000010000010010000101010<br>011000101001110010100000000<br>000110010010001111001101000 | PRESENT FROM | **Acc:** 1.000; **FP:** 0; **FN:** 0 |
| Pool 3 | oLF877/871 | 100000000001111010101010010 | OUR | **Acc:** 1.000; **FP:** 0; **FN:** 0 |

**Extended Data Figure 12: Random access of selected words from the Voyager greeting.** **(a)** Gel electrophoresis of PCR amplicons from Voyager DNA data pools encoded by SpCas9-CBE or SpCas9-ABE with barcode-specific oligonucleotides. (b) Voyager greeting and words selected for random access recovery from SpCas9-CBE encoded DNA data (red circles) or SpCas9-ABE encoded data (purple circles). (c) Random access recovery results. Amplicon pool (Sample), barcode oligonucleotides, binary code after majority vote local clustering decoding of editing rates recorded in amplicons with complete 6-bits in black font, recovered text after binary code conversion, and decoding metrics.

### References


1 Komor, A. C., Kim, Y. B., Packer, M. S., Zuris, J. A. & Liu, D. R. Programmable editing of a target base in genomic DNA without double-stranded DNA cleavage. *Nature* **533**, 420–424 (2016). https://doi.org/10.1038/nature17946

2 Fredens, J. *et al*. Total synthesis of Escherichia coli with a recoded genome. *Nature* **569**, 514–518 (2019). https://doi.org/10.1038/s41586-019-1192-5

**Description of Supplementary Data**

**Supplementary Data 1 – Table of spacer sequences.**
Nucleotide sequences used for spacers in CONCORD experiments.

**Supplementary Data 2 – Table of oligos, plasmids, gBlocks and strains.**
Nucleotide sequences for oligonucleotides and synthesized gene blocks, description of plasmids and genomically engineered bacterial strains.

**Supplementary Data 3 – Trained local and global Thresholds.**

**Supplementary Data 4 – Decoding results for CONCORD training data sets.**

**Supplementary Data 5 – Decoding results for Voyager text and Petroglyph image.**

**Supplementary Data 6 – Programmed binary code and editing rates for CONCORD training data sets.**

**Supplementary Data 7 – Programmed binary code and editing rates for voyager and petroglyph messages.**

**Supplementary Data 8 – Genbank file of an sgRNA plasmid (pLF208) representative of all sgRNA plasmids.**

**Supplementary Data 9 – Genbank file of the SpCas9-CBE base editor plasmid.**

**Supplementary Data 10 – Genbank file of the SpCas9VQR-CBE base editor plasmid.**

**Supplementary Data 11 – Genbank file of the SpCas9EQR-CBE base editor plasmid.**

**Supplementary Data 12 – Genbank file of the SpCas9VRER-CBE base editor plasmid.**

**Supplementary Data 13 – Genbank file of the SpCas9VRQR-CBE base editor plasmid.**

**Supplementary Data 14 – Genbank file of the SpyMacCas9-CBE base editor plasmid.**

**Supplementary Data 15 – Genbank file of the SpCas9-ABE base editor plasmid.**

**Supplementary Data 16 – Genbank file of the SpyMacCas9-ABE base editor plasmid.**

**Supplementary Data 17 – Genbank file of the lacZ locus with integrated synthetic writing sequence for orthogonal base editing testing.**

**Supplementary Data 18 – Genbank file of the lacZ locus with integrated synthetic writing sequence with 32 target positions flanked by NGG PAMs.**

**Supplementary Data 19 – Genbank file of the lacZ locus with integrated synthetic writing sequence with 32 target positions flanked by NAA PAMs.**

**Supplementary Data 20 – Comparison of de novo synthesis-free DNA data storage work**

**Supplementary Data 21 – Custom codes used for denoising, read sampling and data decoding in this study.**
